# Genetic Association Study of Childhood Aggression across raters, instruments and age

**DOI:** 10.1101/854927

**Authors:** Hill F. Ip, Camiel M. van der Laan, Eva M. L. Krapohl, Isabell Brikell, Sánchez-Mora Cristina, Ilja M. Nolte, Beate St Pourcain, Koen Bolhuis, Teemu Palviainen, Hadi Zafarmand, Lucía Colodro-Conde, Scott Gordon, Tetyana Zayats, Fazil Aliev, Chang Jiang, Carol A. Wang, Gretchen Saunders, Ville Karhunen, Anke R. Hammerschlag, Daniel E. Adkins, Richard Border, Roseann E. Peterson, Joseph A. Prinz, Elisabeth Thiering, Ilkka Seppälä, Vilor-Tejedor Natàlia, Tarunveer S. Ahluwalia, Felix R. Day, Jouke-Jan Hottenga, Andrea G. Allegrini, Kaili Rimfeld, Qi Chen, Yi Lu, Joanna Martin, María Soler Artigas, Paula Rovira, Rosa Bosch, Gemma Español, Josep Antoni Ramos Quiroga, Alexander Neumann, Judith Ensink, Katrina Grasby, José J. Morosoli, Xiaoran Tong, Shelby Marrington, Christel Middeldorp, James G. Scott, Anna Vinkhuyzen, Andrey A. Shabalin, Robin Corley, Luke M. Evans, Karen Sugden, Silvia Alemany, Lærke Sass, Rebecca Vinding, Kate Ruth, Jess Tyrrell, Gareth E. Davies, Erik A. Ehli, Fiona A. Hagenbeek, Eveline De Zeeuw, Toos C.E.M. Van Beijsterveldt, Henrik Larsson, Harold Snieder, Frank C. Verhulst, Najaf Amin, Alyce M. Whipp, Tellervo Korhonen, Eero Vuoksimaa, Richard J. Rose, André G. Uitterlinden, Andrew C. Heath, Pamela Madden, Jan Haavik, Jennifer R. Harris, Øyvind Helgeland, Stefan Johansson, Gun Peggy S. Knudsen, Pal Rasmus Njolstad, Qing Lu, Alina Rodriguez, Anjali K. Henders, Abdullah Mamun, Jackob M. Najman, Sandy Brown, Christian Hopfer, Kenneth Krauter, Chandra Reynolds, Andrew Smolen, Michael Stallings, Sally Wadsworth, Tamara L. Wall, Judy L. Silberg, Allison Miller, Liisa Keltikangas-Järvinen, Christian Hakulinen, Laura Pulkki-Råback, Alexandra Havdahl, Per Magnus, Olli T. Raitakari, John R.B. Perry, Sabrina Llop, Maria-Jose Lopez-Espinosa, Klaus Bønnelykke, Hans Bisgaard, Jordi Sunyer, Terho Lehtimäki, Louise Arseneault, Marie Standl, Joachim Heinrich, Joseph Boden, John Pearson, L John Horwood, Martin Kennedy, Richie Poulton, Lindon J. Eaves, Hermine H. Maes, John Hewitt, William E. Copeland, Elizabeth J. Costello, Gail M. Williams, Naomi Wray, Marjo-Riitta Järvelin, Matt McGue, William Iacono, Avshalom Caspi, Terrie E. Moffitt, Andrew Whitehouse, Craig E. Pennell, Kelly L. Klump, S. Alexandra Burt, Danielle M. Dick, Ted Reichborn-Kjennerud, Nicholas G. Martin, Sarah E. Medland, Tanja Vrijkotte, Jaakko Kaprio, Henning Tiemeier, George Davey Smith, Catharina A. Hartman, Albertine J. Oldehinkel, Miquel Casas, Marta Ribasés, Paul Lichtenstein, Sebastian Lundström, Robert Plomin, Meike Bartels, Michel G. Nivard, Dorret I. Boomsma

## Abstract

Childhood aggressive behavior (AGG) has a substantial heritability of around 50%. Here we present a genome-wide association meta-analysis (GWAMA) of childhood AGG, in which all phenotype measures across childhood ages from multiple assessors were included. We analyzed phenotype assessments for a total of 328 935 observations from 87 485 children aged between 1.5 and 18 years, while accounting for sample overlap. We also meta-analyzed within subsets of the data – i.e. within rater, instrument and age. SNP-heritability for the overall meta-analysis (AGG_overall_) was 3.31% (SE=0.0038). We found no genome-wide significant SNPs for AGG_overall_. The gene-based analysis returned three significant genes: *ST3GAL3* (*P*=1.6E-06), *PCDH7* (*P*=2.0E-06) and *IPO13* (*P*=2.5E-06). All three genes have previously been associated with educational traits. Polygenic scores based on our GWAMA significantly predicted aggression in a holdout sample of children (variance explained = 0.44%) and in retrospectively assessed childhood aggression (variance explained = 0.20%). Genetic correlations (*r*_*g*_) among rater-specific assessment of AGG ranged from *r*_*g*_ =0.46 between self- and teacher-assessment to *r*_*g*_ =0.81 between mother- and teacher-assessment. We obtained moderate to strong *r*_*g*_’s with selected phenotypes from multiple domains, but hardly with any of the classical biomarkers thought to be associated with AGG. Significant genetic correlations were observed with most psychiatric and psychological traits (range |*r*_*g*_| : 0.19 – 1.00), except for obsessive-compulsive disorder. Aggression had a negative genetic correlation (*r*_*g*_ =~ −0.5) with cognitive traits and age at first birth. Aggression was strongly genetically correlated with smoking phenotypes (range |*r*_*g*_| : 0.46 – 0.60). The genetic correlations between aggression and psychiatric disorders were weaker for teacher-reported AGG than for mother- and self-reported AGG. The current GWAMA of childhood aggression provides a powerful tool to interrogate the rater-specific genetic etiology of AGG.

## Introduction

There is a variety of phenotypic definitions of aggressive behavior (AGG), from broadly defined externalizing problems to narrow definitions like chronic physical aggression ^1^. Generally any action performed with the intention to harm another organism can be viewed as AGG ^2,3^. AGG is considered a common human behavior ^4^, with people varying in the degree of AGG they exhibit ^5^. Children typically display AGG early in life, after which symptoms tend to diminish ^6,7^, although in some individuals AGG persists into adulthood ^8^. AGG is also part of numerous childhood and adult disorders ^9^, including oppositional defiant disorder (ODD) and conduct disorder (CD)^10^. In its extreme forms, AGG may be considered a disorder by itself – inflicting a huge personal and financial burden on the individual, their relatives, friends, and society as a whole ^11^. In general population studies, AGG is commonly treated as a quantitative trait, and pathological AGG has been argued to be best seen as the extreme end of such a continuum ^12–14^. Childhood AGG co-occurs with many other behavioral, emotional, and social problems ^15,16^ and is associated with increased risk of developing negative outcomes later in life, including cannabis abuse ^17^, criminal convictions ^18^, anxiety disorder ^19^, or antisocial personality disorder ^20^. Not all associated outcomes are harmful ^21^. For example, children who learn to control their impulses and apply aggressive acts as a well-timed coercion strategy are generally more liked by their peers and score higher on social dominance ^22^.

Despite a heritability of roughly 50% ^5,23^, genome-wide association studies (GWASs) on childhood AGG have not identified genome-wide significant loci that replicated ^1^. Childhood cohorts often have rich longitudinal data and assessments from multiple informants and we aimed to increase power to detect genomic loci via multivariate genome-wide association meta-analysis (GWAMA) across genetically correlated traits ^24,25^. In AGG, twin studies have reported moderate to high genetic correlations among instruments, raters, and age [26–29]. Childhood behavior can be context dependent, with teachers, fathers, and mothers each observing and rating aggression against a different background. Teachers are typically unrelated to the child, and see the child in the context of a structured classroom and can judge the child’s behavior against that of other pupils. Parents share part of their genome with their offspring and, most often, a household. Parental genomes also influence the home environment, and it is predominantly within this context that parents observe the child’s behavior. Multiple assessments of aggression by teachers, fathers, and mothers, by different instruments and at different ages, provides information that may be unique to a specific context and therefore may capture context-dependent expression of AGG. These considerations support an approach in which all AGG data are simultaneously analyzed, while retaining the ability to analyze the data by rater. Our analyses include repeated observations on the same subject, which requires appropriate modeling of the clustered data, since the covariance between test statistics becomes a function of a true shared genetic signal and the phenotypic correlation among outcomes ^29^. We developed an approach that allowed inclusion of all measures for a child – e.g. from multiple raters at multiple ages – and resolved issues of sample overlap at the level of the meta-analysis. By doing so we make full use of all data and maximize statistical power for gene discovery. At the same time, by aggregating data at the level of the meta-analysis we retain the flexibility to estimate *r*_*g*_’s between AGG at different ages, by different raters and instruments, and test how AGG assessed by multiple raters differ in the *r*_*g*_ with other phenotypes.

Data on AGG from parent-, teacher- and self-report in boys and girls were collected in 29 cohorts from Europe, USA, Australia, and New-Zealand with 328 935 observations from 87 485 participants, aged 1.5 to 18 years. First, we combined all data to produce the largest GWAMA on childhood AGG to date. SNP-based association tests were followed up by gene-based analyses. We computed polygenic scores (PGSs) to test the out-of-sample prediction of AGG to explore the usefulness of our GWAMA in future research ^30^. To assess genetic pleiotropy between AGG and associated traits, we estimated *r*_*g*_’s with a preselected set of external phenotypes from multiple domains – with a focus on psychiatric and psychological traits, cognition, anthropometric and reproductive traits, substance use, and classic biomarkers of AGG, including testosterone levels. Second, meta-analyses were done by rater, instrument, and age. We estimated *r*_*g*_’s across these assessments of AGG. To identify context-specific genetic overlap with the external phenotypes, *r*_*g*_’s were also estimated between rater-specific assessments of AGG and the external phenotypes.

## Methodology

### Data description

Extended description of the cohorts and phenotypes is supplied in the Supplemental Text and Supplementary Tables 1-9. Cohorts with assessment of AGG in genotyped children and adolescents took part in the meta-analysis. AGG was assessed on continuous scales, with higher scores indicating higher levels of AGG. Within cohort, samples were stratified by (1) rater, (2) instrument and (3) age, maintaining at least 450 observations in each stratum. We ran a univariate GWAS for each stratum within each cohort (Supplementary Table 8). GWASs were run by local analysts following a standard operation protocol (see URLs) after which the summary statistics were uploaded to a central location for the meta-analysis. To account for dependence within cohort in the meta-analysis (see Supplementary Text), each cohort supplied the phenotypic covariance matrix between the AGG measures (Supplementary Table 10) and the degree of sample overlap (Supplementary Table 11) between the different strata. Supplementary Figure 1 shows the distribution of phenotypic correlations across all AGG measures. We assumed no sample overlap across cohorts, and phenotypic correlations among cohorts were set to zero and omitted from Supplementary Figure 1. Phenotypic correlations of zero also correspond to independent samples within a cohort. For GWASs with sample overlap, most phenotypic correlations ranged between 0.1 and 0.4, with a median value of 0.29. When stratified by rater, phenotypic correlations were more heavily centered around 0.4 (see Supplementary Figure 1). The maximum number of correlations within cohort at a specific age is three based on four raters, with the largest number of observations within age-bin around age 12 years. Within this age group, phenotypic correlations among raters ranged between 0.22 and 0.65, with a median of 0.34. The lowest phenotypic correlations were seen between teachers and parents. Since limited data were available on individuals of non-European ancestry, we restricted analyses to individuals of European ancestry.

In total, 29 cohorts contributed 163 GWASs, based on 328 935 observations from 87 485 unique individuals (Supplementary Table 2). Children were 1.5 to 18 years old at assessment, or retrospectively assessed at these ages. Cohorts supplied between 1 and 26 univariate GWASs. Approximately 50% of the subjects were males. Most GWASs were based on maternal- (52.4%) and self-assessment (25.1%), with the remainder based on teacher (12.4%) and paternal report (10.1%). After QC, applied to the univariate GWASs, between 3.47M SNPs and 7.28M SNPs were retained for meta-analysis (see Supplementary Figure 2 and Supplementary Table 9). Note that the wide range of retained SNPs is a result of applying more stringent QC filters for GWASs with smaller sample sizes and that GWASs with comparable sample sizes returned roughly equal number of SNPs (see Supplementary Text and Supplementary Figure 2).

### Meta-analysis

Within cohort measures of AGG may be dependent due to including repeated measures of AGG over age and measures from multiple raters. To account for the effect of sample overlap, we applied a modified version of the multivariate meta-analysis approach developed by Baselmans *et al* ^25^ (see Table 1). Instead of estimating the dependence among GWASs based on the cross-trait-intercept (CTI) with linkage disequilibrium score regression (LDSC)^29,31^, the expected pairwise CTI value was calculated (Table 1) using the observed sample overlap and phenotypic covariance as sample sizes of the univariate GWASs were insufficient to run bivariate LDSC. The effective sample size (N_eff_) was approximated by the third formula in Table 1. When there is no sample overlap (or a phenotypic correlation equal to zero) between all GWASs (i.e. CTI is an identity matrix), N_eff_ is equal to the sum of sample sizes.

**Table 1.** (a) multivariate test statistic in the meta-analysis of results based on overlapping samples. (b) expected value for the cross-trait-intercept. (c) Effective sample size for a GWAMA.

|  |  |
| --- | --- |
| $Z_{multi,j}$ $= \frac{\sum_{i=1}^P w_{ji} Z_{ji}}{\sqrt{\sum_{i=1}^P w_{ji} V_{ji} + \sum_{i=1}^P \sum_{k=1}^P \sqrt{w_{ji} w_{jk}} CTI_{ik} \text{ for } i \neq k}}$ | (a)<br>Multivariate test-statistic for $j$-th SNP. $P$ is the number of GWASs across which we run the meta-analysis; $w_{ji} = \sqrt{N_{ji} h_{SNP,i}^2}$ is the weight given to the $j$th SNP in GWAS $i$, with $h_{SNP,i}^2$ being the SNP-heritability of the trait analyzed in GWAS $i$; and $V_{ji} = 1$ represents the variance of the distribution of $Z_{ji}$ under the null hypothesis of no effect. |
| $CTI_{ik} = \frac{N_s r_p}{\sqrt{N_{ji} N_{jk}}}$ | (b)<br>Cross-trait-intercept between GWAS $i$ and $k$. $N_s$ represents the sample overlap; $r_p$ indicates the phenotypic correlation; $N_{ji}$ and $N_{jk}$ are the sample sizes at SNP $j$ for respectively GWASs $i$ and $k$ |
| $N_{eff} = \sqrt{N}^T CTI^{-1} \sqrt{N}$ | (c)<br>$N$ is an $P$-sized vector of sample sizes, and $CTI$ is the $P \times P$ matrix of cross-trait-intercepts. |

First, we meta-analyzed all available GWASs (AGG_overall_). Second, we meta-analyzed all available data within rater (rater-specific GWAMAs). Third, rater-specific age-bins were created for mother- and self-reported AGG based on the mean ages of the subjects in each GWAS (age-specific GWAMA). To ensure that the age-specific GWAMAs would have sufficient power for subsequent analyses, age-bins were created such that the total *univariate* number of observations (N_obs_) exceeded 15 000 (see Supplementary Text and Supplementary Table 12). For father- and teacher-reported AGG there were insufficient data to run age-specific GWAMAs. Fourth, we performed instrument-specific GWAMAs for (1) the ASEBA scales and (2) for the SDQ, because for these two instruments the total *univariate* N_obs_ was over 15 000.

SNPs that had MAF<0.01, N_eff_<15 000, or were observed in less than two cohorts were removed from further analyses. SNP-heritabilities 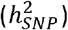 were estimated using LDSC ^31^. *r*_*g*_’s were calculated across stratified assessments of AGG using LDSC ^29^. To ensure sufficient power for the genetic correlations, *r*_*g*_ was calculated across stratified assessments of AGG if the Z-score of the 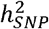 for the corresponding GWAMA was 4 or higher ^29^.

### Gene-based tests

For AGG_overall_, a gene-based analysis was done in MAGMA ^32^. The gene-based test combines P-values from multiple SNPs to obtain a test statistic for each gene, while accounting for LD between the SNPs. From the MAGMA website (see URLs) we obtained (1) a list of 18 087 genes and their start- and end-positions, and (2) pre-formatted European genotypes from 1 000 Genomes phase 3 for the reference LD. We applied a Bonferroni correction for multiple testing at α=0.05/18 087. A lookup for significant results was performed in GWAS Catalog and PhenoScanner (see URLs).

### Polygenic Scores

All data were meta-analyzed twice more, once omitting all data from the Netherlands Twin Register (NTR) and once omitting the Australian data from the Queensland Institute for Medical Research (QIMR,) and the Mater-University of Queensland Study of Pregnancy (MUSP). As the NTR target sample we considered mother-reported AGG at age 7 (*N*=4 491), which represents the largest NTR univariate stratum. In the QIRM participants, we tested whether our childhood AGG PGS predicted adult retrospective assessment of their own CD behavior during adolescence (*N* = 10 706). We allowed for cohort-specific best practice in the polygenic score analysis. In the NTR, we created 16 sets of PGSs in PLINK1.9 ^33^, with P-value thresholds between 1 and 1.0E-05 (see Supplementary Table 13). The remaining SNPs were clumped in PLINK. We applied an *r*^2^-threshold of 0.5 and minimum clumping distance of 250 000 base pair positions ^33^. Age, age^2^, sex, first five ancestry-based principal components, a SNP-array variable, and interaction terms between sex and age, and sex and age^2^ were defined as fixed effects. To account for relatedness, prediction was performed using generalized equation estimation (GEE) as implemented in the “gee” package (version 4.13-19) in R (version 3.5.3). GEE applies a sandwich correction over the standard errors to account for clustering in the data ^34^. To correct for multiple testing, we applied an FDR correction at α=0.05 for 16 tests. QIMR excluded SNPs with low imputation quality (r^2^ = 0.6) and MAF below 1% and selected the most significant independent SNPs using PLINK1.9 ^35^ (criteria linkage disequilibrium r^2^ = 0.1 within windows of 10 MBp). We calculated different PGS for seven P-value thresholds (p<1e-5, p <0.001, p <0.01, p <0.05, p <0.1, p <0.5, and p <1.0) of the GWAS summary statistics. PGS were calculated from the imputed genotype dosages to the 1 000 Genomes (Phase 3 Release 5) reference panel. We fitted linear mixed models, which controlled for relatedness using a Genetic Relatedness Matrix (GRM) and covariates sex, age, two dummy variables for the GWAS array used, and the first five genetic principal components. The parameters of the model were estimated using GCTA 1.9 ^36^ The linear model was as follows:

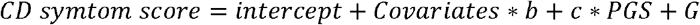

where *b* and *c* represent the vectors of fixed effects; and *G~N*(0,*GRM∗ σ2G*) represents the random effect that models the sample relatedness, with *GRM* being the *N* by *N* matrix of relatedness estimated from SNPs and *N*= 10 706 is the number of individuals.

### Genetic correlations with external phenotypes

We computed *r*_*g*_’s between AGG_overall_ and a set of preselected outcomes (*N*=46; collectively referred to as “external phenotypes”; Supplementary Table 14). Phenotypes were selected based on established hypotheses with AGG and the availability of sufficiently powered GWAS summary statistics. We restricted *r*_*g*_’s to phenotypes for which the Z-scores of the LDSC-based 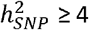 ^29^. Next, we estimated *r*_*g*_’s for all rater-specific assessments of AGG (except for father-reported AGG). Genomic Structural Equation Modelling (Genomic SEM)^37^ was applied to test if *r*_*g*_’s were significantly different across raters. Specifically, for every phenotype, we tested whether (1) all three *r*_*g*_’s between the external phenotype and rater-specific assessment of AGG, i.e. mother, teacher or self-ratings, could be constrained at zero, and (2) whether *r*_*g*_’s could be constrained to be equal across raters. A *χ*^2^ difference test was applied to assess whether imposing the constraints resulted in a significant worse model fit compared to a model where the *r*_*g*_’s between the phenotype and three rater-specific assessment of AGG were allowed to differ. We applied an FDR correction at α=0.05 over two models for 46 external phenotypes, for a total of 92 tests. An FDR correction for 4 x 46=184 tests was applied to correct for multiple testing of whether the genetic correlations were significantly different from zero.

## Results

### Overall GWAMA

We first meta-analyzed the effect of each SNP across all available univariate GWASs. Assuming an N_eff_ of 151 741, the 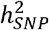 of AGG_overall_ was estimated at 3.31% (SE=0.0038). The mean *χ*^2^-statistic was 1.12 along with an LDSC-intercept of 1.02 (SE=0.01). This indicated that a small, but significant, part of the inflation in test statistics might have been due to confounding biases, which can either reflect population stratification or subtle misspecification of sample overlap within cohorts. No genome-wide significant hits were found for AGG_overall_ (Figure 1). The list of suggestive associations (*P*<1.0E-05) is provided in Supplementary Table 15. SNPs were annotated with SNPnexus (see URLs). The strongest association, in terms of significance, was located on chromosome 2 (rs2570485; *P*=2.0E-07). The SNP is located inside a gene desert, without any gene in 400Kbp in any direction. The second strongest independent association was found with rs113599846 (*P*=4.3E-07), which is located inside an intronic region of *TNRC18* on chromosome 7. None of the suggestive associations have previously been reported for AGG or AGG-related traits ^1^.

**Figure 1.**
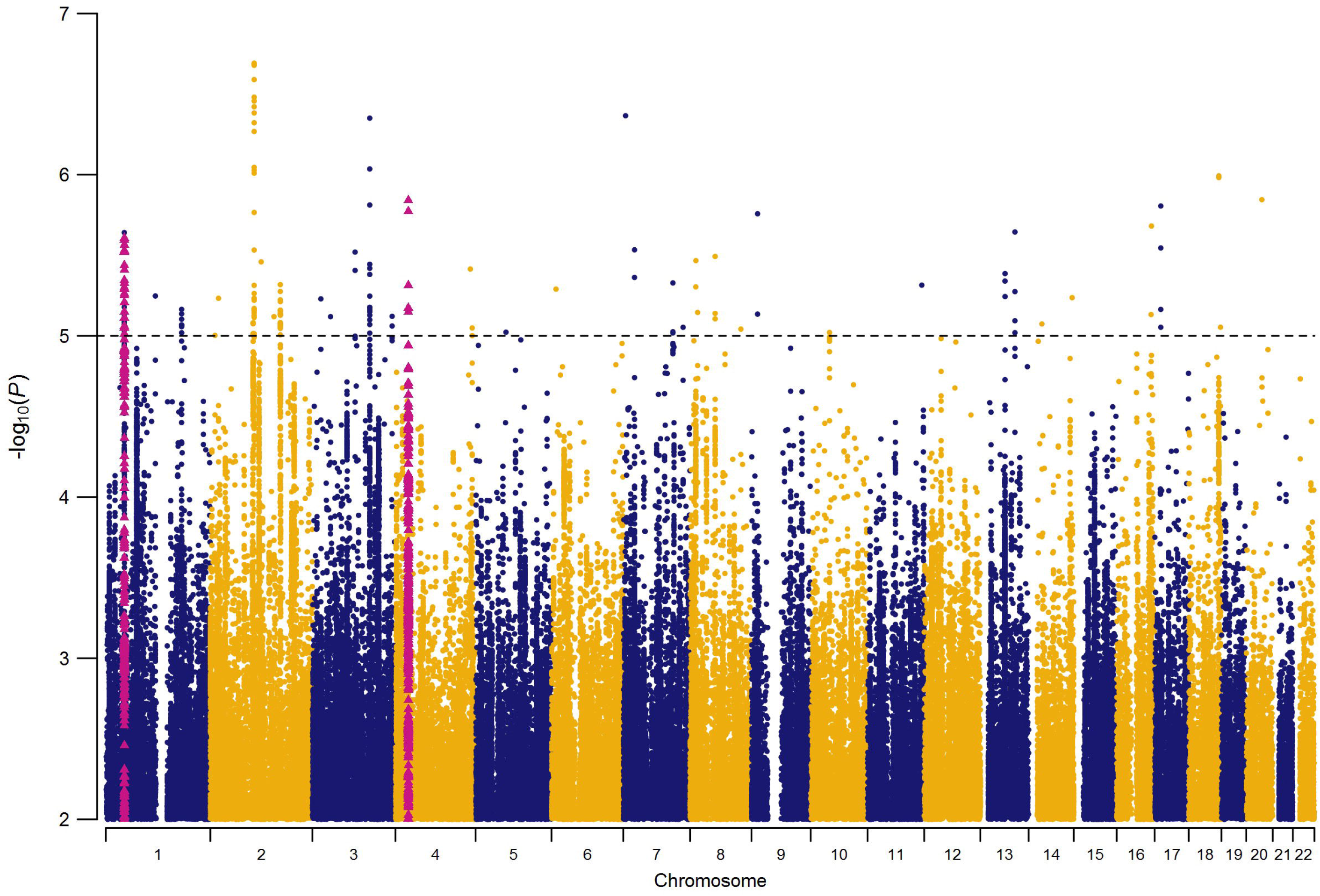
Manhattan plot of overall meta-analysis for childhood aggression (AGG_overall_). Red triangles represent SNPs that were included in the significant genes from the gene-based analysis. SNPs for *ST3GAL3* and *IPO13* are included in the same locus on chromosome 1.

We tested previously reported genome-wide significant associations for AGG ^1^ and performed a lookup in AGG_overall_. We restricted lookup to associations with autosomal SNPs that were found in samples of European ancestry, resulting in three loci. One genome-wide significant hit was reported for adult antisocial personality disorder (rs4714329; OR=0.63^1^; *P*=1.64E-09)^38^. The same SNP, however, had an opposite direction of effect in AGG_overall_ (β=0.0022; *P*=0.41). Tielbeek *et al* ^39^ reported two genome-wide significant hits for antisocial behavior, one on chromosome 1 (rs2764450) and one on chromosome 11 (rs11215217). While both SNPs have the same direction of effect, neither SNP is associated with AGG_overall_ (both *P*>0.5).

### Gene-based analysis

After correction for multiple testing, the gene-based analysis returned three significant results (Supplementary Table 16): *ST3GAL3* (ST3 beta-galactoside alpha-2,3-sialyltransferase3; *P*=1.6E-06), *PCDH7* (protocadherin 7; *P*=2.0E-06) and *IPO13* (importin 13; *P*=2.5E-06). *ST3GAL3* codes for a type II membrane protein that is involved in catalyzing the transfer of sialic acid from CMP-sialic acid to galactose-containing substrates. *ST3GAL3* has been implicated in 107 GWASs, most notably on intelligence and educational attainment. The top SNP in *ST3GAL3* (rs2485997; *P*=2.48E-06) is in strong LD (r^2^>0.8) with several other SNPs inside the gene body of *ST3GAL3* and in moderate LD (r^2^>0.6) with SNPs in several neighboring genes (Supplementary Figure 3). *PCDH7* codes for a protein that is hypothesized to function in cell-cell recognition and adhesion. *PCDH7* has been implicated in 196 previous GWASs, for example educational attainment and adventurousness. The top SNP for *PCDH7* (rs13138213; *P*=1.44E-06) is in strong LD (r^2^>0.8) with a small number of other closely located SNPs and the signal for the gene-based test appears to be driven by two independent loci (Supplementary Figure 4). *IPO13* codes for a nuclear transport protein. *IPO13* has been implicated in the UKB GWASs on whether a person holds a college or university degree and intelligence. The top SNP (rs3791116; *P*=1.19E-05) is in moderate to strong LD with multiple SNPs (Supplementary Figure 5), including SNPs in the neighboring *ST3GAL3* gene.

### Polygenic prediction

In children, 11 out of 16 polygenic scores were significantly correlated with mother-reported AGG in 7-year-olds (Figure 2) after correction for multiple testing. The scores explained between 0.036% and 0.44% of the phenotypic variance. The significant correlations consistently emerged when scores including SNPs with P-values above 0.002 in the discovery GWAS were considered. In the retrospective assessments of adolescent CD, the PGS calculated at various thresholds (Figure 3) explained up to 0.2% of the variance in symptom sum scores. Generally, CD is significantly predicted at most thresholds, although, as we would expect based on the SNP-heritability of AGG_overall_, the proportion of explained variance is small.

**Figure 2.**
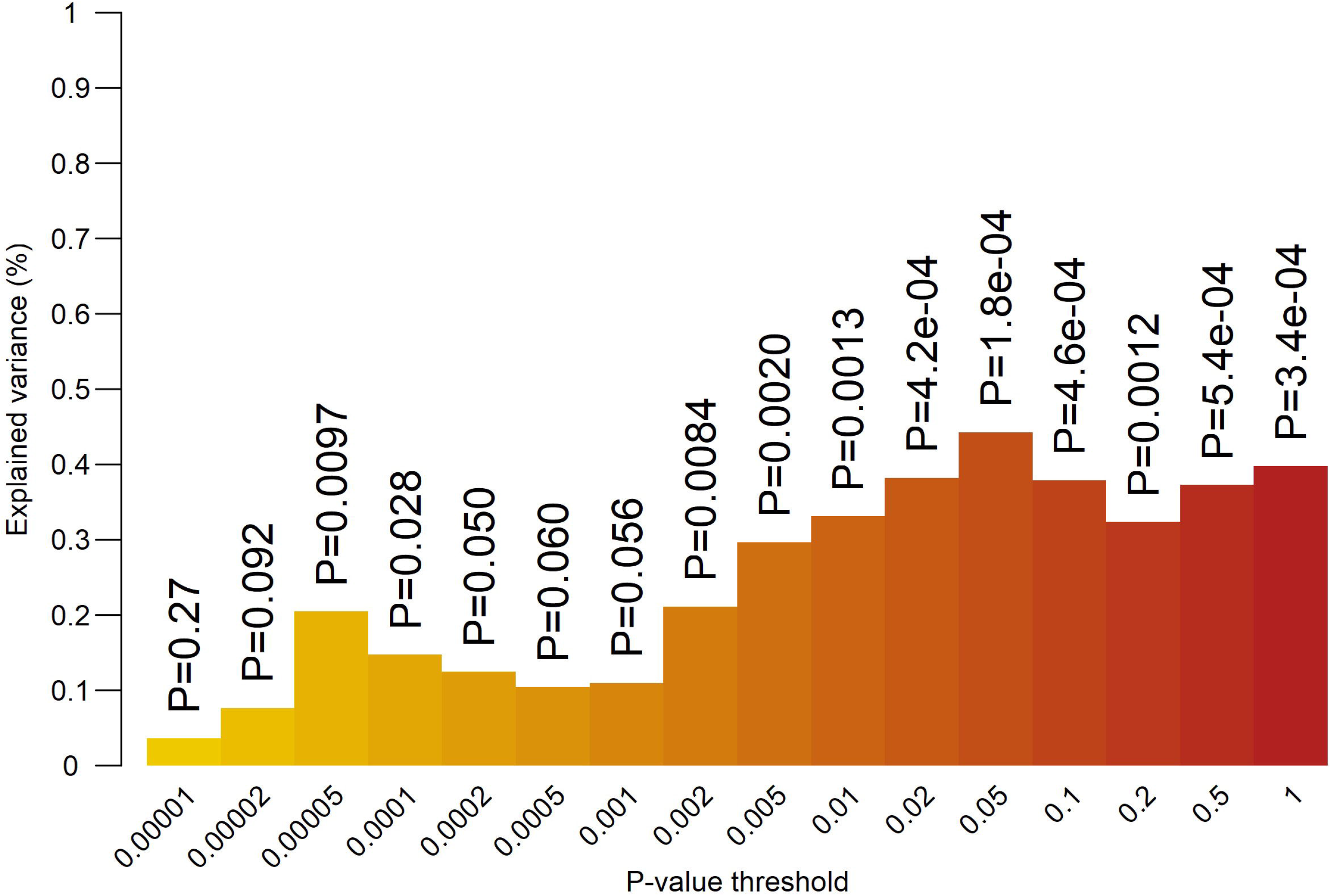
Proportion of explained variance (vertical axis) in childhood aggression at age 7 by polygenic scores from the overall GWAMA for multiple *P*-value thresholds (horizontal axis). Numbers above the bars represent unadjusted *P*-values for two-sided test of significance.

**Figure 3.**
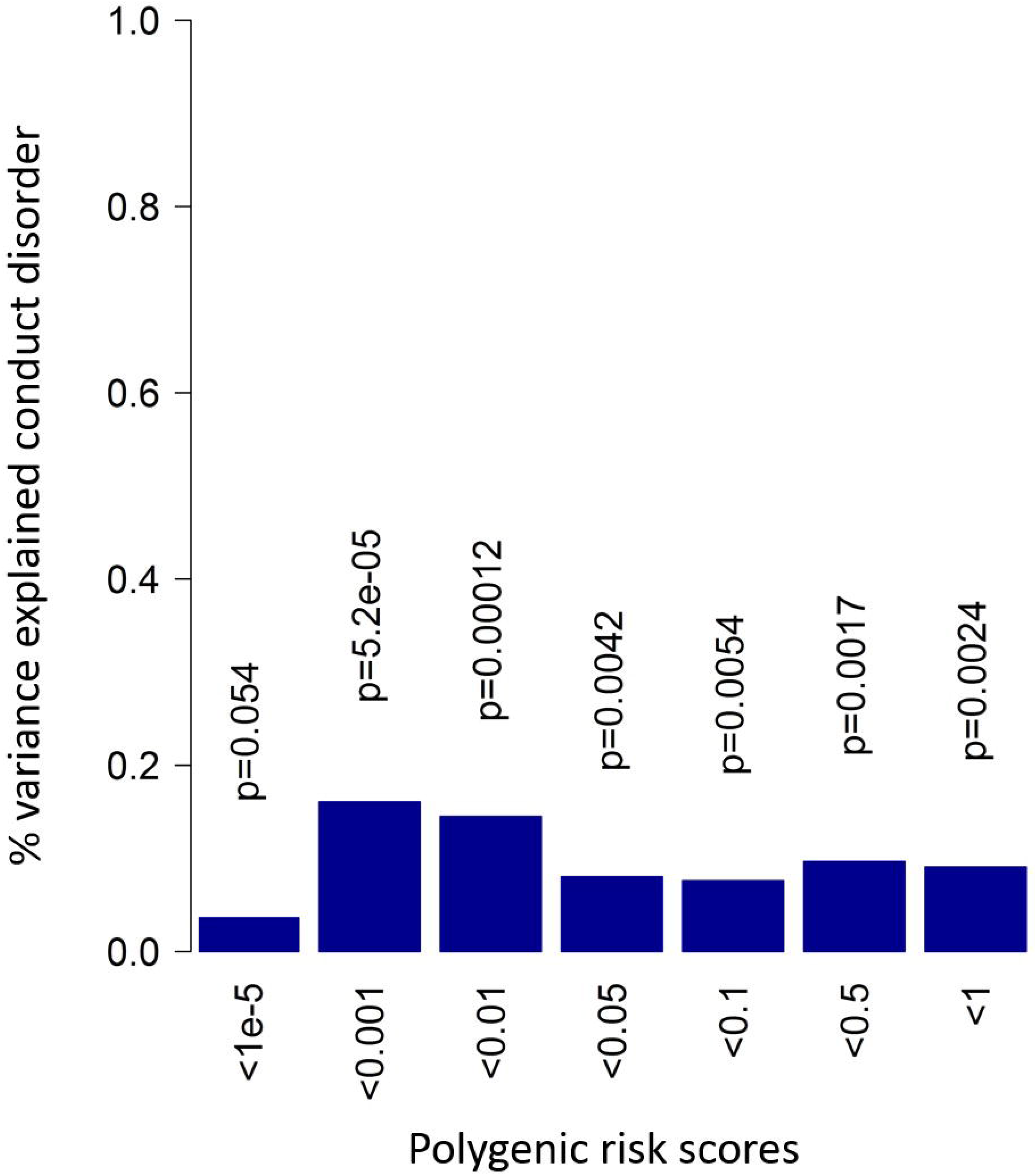
Proportion of explained variance (vertical axis) in retrospective adolescent CD (two sided tests). Blue bars indicate positive correlation with the conduct disorder score.

### Genetic correlation with external phenotypes

Genetic correlations between AGG_overall_ and a set of preselected external phenotypes are shown in Figure 4 and Supplementary Table 17. These phenotypes can broadly be grouped into psychiatric and psychological traits, substance use, cognitive ability, anthropometric traits, classic biomarkers of AGG, reproductive traits, and sleeping behavior. We included childhood phenotypes (e.g. birth weight and childhood IQ) and disorders (e.g. ADHD and autism spectrum disorder [ASD]), but the majority of phenotypes were adult characteristics or characteristics measured in adult samples. After correction for multiple testing, 36 phenotypes showed a significant *r*_*g*_ with AGG_overall_ (*P*<0.02). In general, the highest positive correlations were seen with psychiatric traits, notably ADHD, ASD, and major depressive disorder (MDD). The largest negative genetic correlations were found for age at smoking initiation, childhood IQ, and age at first birth. Based on the biomarker-aggression literature, we tested for the presence of genetic correlations between AGG_overall,_ and lipids, heart rate, heart rate variability, and testosterone levels. Very low genetic correlations were observed for AGG_overall,_ and these biomarkers, with in many cases the sign of the genetic correlation opposite to what was expected based on the literature on biomarkers of AGG.

**Figure 4.**
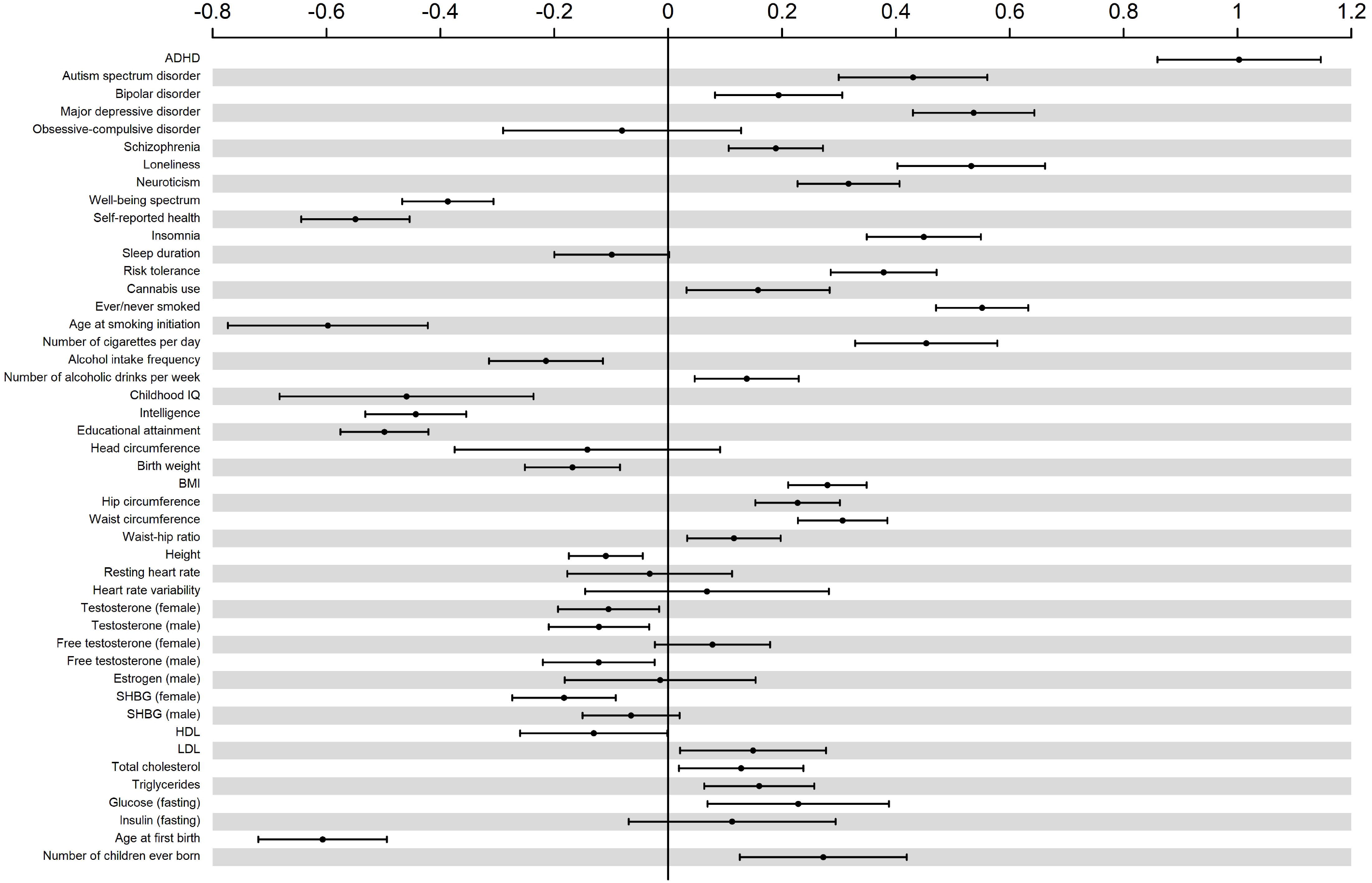
Genetic correlation with external phenotypes. Phenotypes are ordered by domain. Bars indicate 95% confidence intervals.

### Stratified assessment of childhood aggressive behavior

Separate meta-analyses were carried out for raters, instruments and age. None of these GWAMAs returned genome-wide significant hits. Manhattan plots for the four rater-specific GWAMAs are shown in Supplementary Figure 6. Estimates of 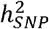 for rater-specific assessment of AGG are shown in Supplementary Table 18. The lowest 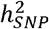 was observed for father-reported AGG (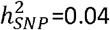; SE=0.03) and the highest for teacher-reported AGG (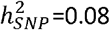; SE=0.02). We estimated *r*_*g*_ between rater-specific assessment of AGG, except for father-reported AGG, which returned a non-significant 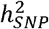. A substantial genetic correlation was observed between AGG_Mother_ and AGG_Teacher_ (*r*_*g*_ =0.81; SE=0.11). Moderate genetic correlations were observed between AGG_Self_ and AGG_Mother_ (*r*_*g*_ =0.67; SE=0.10), and between AGG_Self_ and AGG_Teacher_ (*r*_*g*_ =0.46; SE=0.13). Both genetic correlations involving self-reported AGG were significantly lower than 1.

We performed a GWAMA across all GWASs where an ASEBA scale was used (AGG_ASEBA_) and another GWAMA across all GWASs for the SDQ (AGG_SDQ_). SNP-heritabilities for AGG_ASEBA_ and AGG_SDQ_ were 0.031 (SE=0.0099) and 0.026 (SE=0.0086), respectively. The GWAMAs were insufficiently powered to estimate *r*_*g*_ across instrument-specific assessment of AGG.

Age-specific GWAMAs were performed for mother- and self-reported AGG, which made up 77.5% of the data. Mother-reported data were split into seven age-bins and self-reported data into three (Supplementary Table 12). Estimates of the 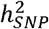 for each age-specific GWAMA can be found in Supplementary Table 19. For mother-reported AGG, 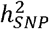 ranged between 0.012 and 0.078. For self-reported AGG, the highest 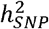 was seen for the retrospective data (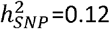; SE=0.03), which also showed a significantly inflated intercept (1.05; SE=0.01). *r*_*g*_ could only be estimated between AGG_M7_, AGG_S13_ and AGG_SR_ (Supplementary Table 20).

### Genetic correlation between rater-specific assessment of AGG and external phenotypes

We estimated rater-specific *r*_*g*_’s with the external phenotypes, except for father-reported AGG, and tested for each external phenotype whether these *r*_*g*_’s could be constrained to be equal to zero. For 31 out of 46 external phenotypes, constraining the *r*_*g*_’s to be equal to zero for all three raters resulted in significant reduction in model fit (Supplementary Table 21), indicating that, for these external phenotypes, at least one rater has an *r*_*g*_ that is significantly different from zero.

Next, we tested for each external phenotype whether the three rater-specific *r*_*g*_’s with the external phenotypes could be constrained to be equal across mothers, teachers and self-ratings. For ADHD, ASD, MDD, schizophrenia, well-being, and self-reported health, constraining the *r*_*g*_’s to be equal across rater resulted in significantly worse model fit (Supplementary Table 21). For all these phenotypes, *r*_*g*_’s with teacher-reported AGG were consistently lower compared to mother- and self-reported AGG (Supplementary Figure 7 and Supplementary Table 17). For lifetime cannabis use, genetic correlations also could not be constrained to be equal across raters. Here, a relatively strong *r*_*g*_ was found with self-reported AGG (*r*_*g*_ =0.36; SE=0.08) compared to teacher- (*r*_*g*_ =0.13; SE=0.07) and mother-reported AGG (*r*_*g*_ =0.08; SE=0.08).

## Discussion

We present the largest genome-wide association meta-analysis (GWAMA) of childhood aggressive behavior (AGG) to date. The gene-based analysis implicated three genes, *PCDH7*, *ST3GAL3* and *IPO13*, based on the overall meta-analysis (AGG_overall_), which did not return genome-wide significant SNPs. Lead SNPs in the implicated genes were related to educational outcomes, but did not reach genome-wide significance and these loci require further evidence before being considered as AGG risk variants. Polygenic scores (PGS) predicted childhood AGG and retrospectively assessed adolescent CD. Stratified analyses within AGG generally returned moderate to strong genetic correlations across raters. We found substantial genetic correlations between AGG_overall_ and a list of preselected external phenotypes from various domains, including, psychiatry and psychology, cognition, anthropometric and reproductive traits. Most notably was the perfect *r*_*g*_ between AGG_overall_ and ADHD (*r*_*g*_ =1.00; SE=0.07). This is in line with the moderate-to-strong phenotypic correlations that have consistently been found across sex-, rater-, age- and instrument-specific assessment of AGG with attention problems and hyperactivity ^15^. Significant genetic correlations were further observed with other psychiatric and psychological traits (range |*r*_*g*_| : 0.19 – 0.55). Negative genetic correlations (*r*_*g*_ =~ −0.5) were found with all three traits from the cognitive domain. Genetic correlations were positive with smoking initiation (*r*_*g*_ =0.55; SE=0.04) and smoking quantity (*r*_*g*_ =0.46; SE=0.06), and negative with age at smoking initiation (*r*_*g*_ =−0.60; SE=0.09).

We examined genetic correlations with classical biomarkers of aggressive behavior. Higher levels of aggression have been associated with lower levels of LDL ^40^ and lower resting heart rate ^41,42^. We found a positive, albeit weak, *r*_*g*_ between AGG_overall_ and LDL (*r*_*g*_ =0.15; SE=0.07), which has an opposite sign than what was expected based on the literature [39]. More broadly, except for HDL (*r*_*g*_ =−0.13; SE=0.07), all measures of lipid levels returned significant positive *r*_*g*_’s with AGG_overall_, albeit weakly (*r*_*g*_ <0.2). No heart rate measure showed a significant genetic correlation with AGG_overall_. The relationship between testosterone levels and (childhood) AGG in the literature is, at best, unclear. A positive association between AGG and testosterone is often assumed, but the relation may be more complex ^43^. Both positive and negative phenotypic correlations have been found and seem context-dependent ^44^. We found significant negative, *r*_*g*_’s between AGG and testosterone levels in males and females (|*r*_*g*_| <0.15). These should be interpreted with some caution because of the design of the GWA studies: AGG was measured in children and young adolescents whereas testosterone levels were measured in adults in the UK Biobank ^45^, and genetic stability of testosterone levels might be low, at least for males ^46^. Genetic correlations with reproductive traits showed a positive relation with having more children (*r*_*g*_ =0.27; SE=0.08) and having offspring earlier in life (*r*_*g*_ =−0.60; SE=0.06), tending to confirm that not all associated outcomes are harmful.

The stratified design of our study also allowed for examination of the genetic etiology of AGG in subsets of the data and examination of genetic correlations among raters. We found a high genetic correlation between AGG_Mother_ and AGG_Teacher_ (*r*_*g*_ =0.81; SE=0.11). However, the 95% confidence interval covers 1, which makes these results hard to reconcile with previous findings of rater-specific additive genetic effects in childhood AGG ^47^. Most external phenotypes showed comparable *r*_*g*_’s with mother-, self-, and teacher-reported AGG. For ADHD, ASD, MDD, schizophrenia, well-being, and self-reported health, *r*_*g*_’s differed significantly across raters. Weaker *r*_*g*_’s were consistently found in teacher-reported AGG compared to mother- and self-reported AGG. These findings indicate the presence of rater-specific effects when considering the genetic correlation of AGG with other outcomes. *r*_*g*_’s are generally stronger in the psychopathology and psychological domains. A lack of power, however, seems insufficient to explain why we found weaker *r*_*g*_’s between AGG_Teacher_ and phenotypes from these two domains. Other phenotypes, like smoking behavior, educational attainment or age at first birth, are, like psychopathological phenotypes, highly genetically correlated with AGG_overall_, but, unlike psychopathologies, have near identical *r*_*g*_’s across raters. The rater-specific effects on *r*_*g*_’s between childhood AGG and external phenotypes might be limited to psychopathologies, and future research into the genetics of childhood psychopathology might consider these nuances in effects of assessment of childhood AGG from various sources, be that multiple raters, instruments, and ages.

Despite the considerable sample sizes, we were still underpowered to compute genetic correlations with external phenotypes while stratifying AGG over age or instrument. Age-stratified GWASs in larger samples across development are a desirable target for future research. Because genetic correlations can be computed between phenotypes for which a well-powered GWAS is available, age-stratified GWAS of many developmental phenotypes, behavioral, cognitive and neuroscientific can be leveraged to better understand development of childhood traits.

We note that multivariate results should be interpreted with some caution. While combining data from correlated traits can indeed improve power to identify genome-wide associations, interpreting the phenotype may not be straightforward. In the current GWAMA, we have referred to our phenotype as “aggressive behavior” and interpreted the results accordingly. Aggressive behavior, however, is an umbrella term that has been used to identify a wide range of distinct – though correlated – traits and behaviors ^1^.

Genome-wide association studies are increasingly successful in identifying genomic loci for complex human traits ^48^ and also in psychiatry, genetic biomarkers are increasingly thought of as promising for both research and treatment. Genetic risk prediction holds promise for adult psychiatric disorders ^30^ and it seems reasonable to expect the same for childhood disorders. Here we found that polygenic scores explain up to 0.44% of the phenotypic variance in AGG in 7-year-olds and 0.2% of the variance in retrospectively reported adolescent CD. Note that differences in ages, instrument and local best-practices have led to differences in explained variance. Future studies may explore the utility of these PGSs in illuminating pleiotropy between AGG_overall_ and other traits. A limiting factor in this regard is the relatively low SNP-heritability, which puts an upper bound on the predictive accuracy of PGSs. Since measurement error suppresses SNP-heritability, better measurement may offer an avenue to higher powered GWAS, and subsequently to better PGS. Furthermore, sample sizes for developmental phenotypes, including AGG may need to increase by one to two orders of magnitude before PGS become useful for individual patients.

Despite our extensive effort, the first genome-wide significant SNP for childhood AGG has yet to be found. Even in the absence of genome-wide significant loci, however, GWASs aid in clarifying the biology behind complex traits. Our results show that, even without genome-wide significant hits, a GWAS can be powerful enough to illuminate the genetic etiology of a trait in the form of *r*_*g*_’s with other complex traits. Non-significant associations are expected to capture part of the polygenicity of a trait ^31^ and various follow up-analyses have been developed for GWASs that do not require, but are aided by, genome-wide significant hits ^49^. Polygenic scores aggregate SNP effects into a weighted sum that indicates a person’s genetic liability to develop a disorder. While their clinical application is still limited in psychiatric disorders, they can already aid in understanding the pleiotropy among psychiatric and other traits ^30^. Similarly, summary statistics-based genetic correlations (*r*_*g*_) provide insight into the genetic overlap between complex traits ^29,50^.

## URLs

GWAS SOP: http://www.action-euproject.eu/content/data-protocols

MAGMA: https://ctg.cncr.nl/software/magma

SNPnexus: https://www.snp-nexus.org/index.html (accessed on 28-8-2019)

GWAS Catalog: https://www.ebi.ac.uk/gwas/ (accessed on 29-8-2019)

PhenoScanner: http://www.phenoscanner.medschl.cam.ac.uk/ (accessed on 29-8-2019)

## Acknowledgements

We very warmly thank all participants, their parents and teachers for making this study possible. The project was supported by the “Aggression in Children: Unraveling gene-environment interplay to inform Treatment and InterventiON strategies” project (ACTION). ACTION received funding from the European Union Seventh Framework Program (FP7/2007-2013) under grant agreement no 602768. Cohort specific acknowledgements and funding information may be found in Supplemental Text.

## Author contributions

may be found in Supplemental Text

## Conflict of interests

Miquel Casas has received travel grants and research support from Eli Lilly and Co., Janssen-Cilag, Shire and Lundbeck and served as consultant for Eli Lilly and Co., Janssen-Cilag, Shire and Lundbeck. Josep Antoni Ramos Quiroga was on the speakers’ bureau and/or acted as consultant Eli-Lilly, Janssen-Cilag, Novartis, Shire, Lundbeck, Almirall, Braingaze, Sincrolab, Medicine, Exeltis and Rubió in the last 5 years. He also received travel awards (air tickets + hotel) for taking part in psychiatric meetings from Janssen-Cilag, Rubió, Shire, Medice and Eli-Lilly. The Department of Psychiatry chaired by him received unrestricted educational and research support from the following companies in the last 5 years: Eli-Lilly, Lundbeck, Janssen-Cilag, Actelion, Shire, Ferrer, Oryzon, Roche, Psious, and Rubió.

## Footnotes

1 odds ratio was signed to the other allele in the original study

## Notes

### Summary of Updates

Updated manuscript

## References

1 Odintsova V V et al. Genomics of human aggression: current state of genome-wide studies and an automated systematic review tool. Psychiatr Genet 2019; 29: 170–190.

2 Baron RA, Richardson DR. Human Aggression. 2nd ed. New York, 1994.

3 Anderson CA, Bushman BJ. Human aggression. Annu Rev Psychol 2002; 53: 27–51.

4 Veroude K et al. Genetics of aggressive behavior: An overview. Am J Med Genet Part B Neuropsychiatr Genet 2016; 171: 3–43.

5 Tuvblad C, Baker LA. Human aggression across the lifespan: genetic propensities and environmental moderators. Adv Genet 2011; 75: 171–214.

6 Bongers IL, Koot HM, van der Ende J, Verhulst FC. Developmental Trajectories of Externalizing Behaviors in Childhood and Adolescence. Child Dev 2004; 75: 1523–1537.

7 Tremblay RE, Vitaro F, Côté SM. Developmental Origins of Chronic Physical Aggression: A Bio-Psycho-Social Model for the Next Generation of Preventive Interventions. Annu Rev Psychol 2018; 69: 383–407.

8 Huesmann LR, Dubow EF, Boxer P. Continuity of aggression from childhood to early adulthood as a predictor of life outcomes: Implications for the adolescent-limited and life-course-persistent models. Aggress Behav 2009; 35: 136–149.

9 Zhang-James Y, Faraone S V. Genetic architecture for human aggression: A study of gene-phenotype relationship in OMIM. Am J Med Genet Part B Neuropsychiatr Genet 2016; 171: 641–649.

10 American Psychiatric Association. Diagnostic and Statistical Manual of Mental Disorders, Fifth Edition. American Psychiatric Association: Arlington, 2013.

11 Foster EM, Jones DE, The Conduct Problems Prevention Research Group. The high costs of aggression: public expenditures resulting from conduct disorder. Am J Public Health 2005; 95: 1767–72.

12 Walton KE, Ormel J, Krueger RF. The dimensional nature of externalizing behaviors in adolescence: Evidence from a direct comparison of categorical, dimensional, and hybrid models. J Abnorm Child Psychol 2011; 39: 553–561.

13 Walters GD, Ruscio J. Trajectories of youthful antisocial behavior: Categories or continua? J Abnorm Child Psychol 2013; 41: 653–666.

14 Barry TD, Marcus DK, Barry CT, Coccaro EF. The latent structure of oppositional defiant disorder in children andadults. J Psychiatr Res 2013; 47: 1932–1939.

15 Bartels M et al. Childhood aggression and the co-occurrence of behavioural and emotional problems: results across ages 3–16 years from multiple raters in six cohorts in the EU-ACTION project. Eur Child Adolesc Psychiatry 2018; 27: 1105–1121.

16 Whipp AM et al. Teacher-rated aggression and co-occurring problems and behaviors among schoolchildren: A comparison of four population-based European cohorts. medRxiv 2019; : 19002576.

17 Heron J et al. Childhood conduct disorder trajectories, prior risk factors and cannabis use at age 16: birth cohort study. Addiction 2013; 108: 2129–2138.

18 Rivenbark JG et al. The high societal costs of childhood conduct problems: evidence from administrative records up to age 38 in a longitudinal birth cohort. J Child Psychol Psychiatry 2018; 59: 703–710.

19 Odgers CL et al. Female and male antisocial trajectories: From childhood origins to adult outcomes. Dev Psychopathol 2008; 20: 673–716.

20 Whipp AM et al. Early adolescent aggression predicts antisocial personality disorder in young adults: a population-based study. Eur Child Adolesc Psychiatry 2019; 28: 341–350.

21 Hawley PH, Little TD, Rodkin PC. Aggression and adaptaion: The bright side to bad behavior. 1st ed. Routledge: New York, 2007 doi:https://doi.org/10.4324/9780203936900.

22 Hawley PH. Social Dominance in Childhood and Adolescence: Why Social Competence and Aggression May Go Hand in Hand. In: Hawley PH, Little TD, Rodkin PC (eds). Aggression and Adaptation: The Bright Side to Bad Behavior. Routledge: New York, 2007, pp 1–22.

23 Waltes R, Chiocchetti AG, Freitag CM. The neurobiological basis of human aggression: A review on genetic and epigenetic mechanisms. Am J Med Genet Part B Neuropsychiatr Genet 2016; 171: 650–675.

24 Turley P et al. Multi-trait analysis of genome-wide association summary statistics using MTAG. Nat Genet 2018; 50: 229–237.

25 Baselmans BML et al. Multivariate genome-wide analyses of the well-being spectrum. Nat Genet 2019; : 1.

26 van Beijsterveldt CEM, Bartels M, Hudziak JJ, Boomsma DI. Causes of Stability of Aggression from Early Childhood to Adolescence: A Longitudinal Genetic Analysis in Dutch Twins. Behav Genet 2003; 33: 591–605.

27 Porsch RM et al. Longitudinal heritability of childhood aggression. Am J Med Genet Part B Neuropsychiatr Genet 2016; 171: 697–707.

28 Lubke GH, McArtor DB, Boomsma DI, Bartels M. Genetic and environmental contributions to the development of childhood aggression. Dev Psychol 2018; 54: 39–50.

29 Bulik-Sullivan BK et al. An atlas of genetic correlations across human diseases and traits. Nat Genet 2015; 47: 1236–1241.

30 Martin AR, Daly MJ, Robinson EB, Hyman SE, Neale BM. Predicting Polygenic Risk of Psychiatric Disorders. Biol Psychiatry 2019; 86: 97–109.

31 Bulik-Sullivan BK et al. LD Score regression distinguishes confounding from polygenicity in genome-wide association studies. Nat Genet 2015; 47: 291–295.

32 de Leeuw CA, Mooij JM, Heskes T, Posthuma D. MAGMA: Generalized Gene-Set Analysis of GWAS Data. PLOS Comput Biol 2015; 11: e1004219.

33 Chang CC et al. Second-generation PLINK: rising to the challenge of larger and richer datasets. Gigascience 2015; 4: 7.

34 Rogers P, Stoner J. Modification of the Sandwich Estimator in Generalized Estimating Equations with Correlated Binary Outcomes in Rare Event and Small Sample Settings. Am J Appl Math Stat 2015; 3: 243–251.

35 Purcell S et al. PLINK: a tool set for whole-genome association and population-based linkage analyses. Am J Hum Genet 2007; 81: 559–575.

36 Yang J, Lee SH, Goddard ME, Visscher PM. GCTA: a tool for genome-wide complex trait analysis. Am J Hum Genet 2011; 88: 76–82.

37 Grotzinger AD et al. Genomic structural equation modelling provides insights into the multivariate genetic architecture of complex traits. Nat Hum Behav 2019; : 1.

38 Rautiainen M-R et al. Genome-wide association study of antisocial personality disorder. Transl Psychiatry 2016; 6: e883–e883.

39 Tielbeek JJ et al. Genome-Wide Association Studies of a Broad Spectrum of Antisocial Behavior. JAMA Psychiatry 2017; 74: 1242.

40 Hagenbeek FA et al. Adult aggressive behavior in humans and biomarkers: a focus on lipids and methylation. J Pediatr Neonatal Individ Med 2018; 7: e070204.

41 Raine A, Fung ALC, Portnoy J, Choy O, Spring VL. Low heart rate as a risk factor for child and adolescent proactive aggressive and impulsive psychopathic behavior. Aggress Behav 2014; 40: 290–299.

42 Latvala A et al. Association of Resting Heart Rate and Blood Pressure in Late Adolescence With Subsequent Mental Disorders. JAMA Psychiatry 2016; 73: 1268.

43 Brain PF, Susman EJ. Hormonal aspects of aggression and violence. In: Handbook of antisocial behavior. John Wiley & Sons Inc.: Hoboken, 1997, pp 314–323.

44 Ramirez JM. Hormones and aggression in childhood and adolescence. Aggress Violent Behav 2003; 8: 621–644.

45 Bycroft C et al. The UK Biobank resource with deep phenotyping and genomic data. Nature 2018; 562: 203–209.

46 Harris JA, Vernon PA, Boomsma DI. The Heritability of Testosterone: A Study of Dutch Adolescent Twins and Their Parents. Behav Genet 1998; 28: 165–171.

47 Hudziak JJ et al. Individual Differences in Aggression: Genetic Analyses by Age, Gender, and Informant in 3-, 7-, and 10-Year-Old Dutch Twins. Behav Genet 2003; 33: 575–589.

48 Visscher PM et al. 10 years of GWAS discovery: biology, function, and translation. Am J Hum Genet 2017; 101: 5–22.

49 Pasaniuc B, Price AL. Dissecting the genetics of complex traits using summary association statistics. Nat Rev Genet 2017; 18: 117–127.

50 Watanabe K et al. A global overview of pleiotropy and genetic architecture in complex traits. Nat Genet 2019; : 1–10.

