## Supplementary Text for "Genetic Association Study of Childhood Aggression across raters, instruments and age"

**Cohort description**

*ABCD*

Data for this study comes from the ABCD-Genetic Enrichment (ABCD-GE) study, a sub-study of 1 192 ethnic Dutch children. Approval of the study was obtained from the Central Committee on Research Involving Human Subjects in the Netherlands, the medical ethics review committees of the participating hospitals, and the Registration Committee of the Municipality of Amsterdam and written consent was obtained from participating parents and children of the phenotypes. Regarding the DNA collection and analysis, an opt-out procedure was used (METC approval 2002_039#B2013531).

*ALSPAC*

Data for this study were obtained for children of the ALSPAC study, a UK population-based longitudinal pregnancy-ascertained birth cohort (estimated birth date: 1991-1992)[1, 2]. Ethical approval was obtained from the ALSPAC Law-and-Ethics Committee (IRB00003312) and the Local Research-Ethics Committees. Written informed consent was obtained from a parent or individual with parental responsibility and assent (and for older children consent) was obtained from the child participants. The study website contains details of all the data that is available through a fully searchable data dictionary (<http://www.bris.ac.uk/alspac/researchers/data-access/data-dictionary/>).

*BREATHE*

Participants were drawn from the BREATHE project (European Commission: FP7-ERC-2010-AdG, ID 268479), a population-based cohort of primary schoolchildren designed to analyze the association between air pollution and behavior, cognitive function and brain morphology [3]. A total of 2 897 children aged 7 to 11 years accepted the invitation and participated in the project. Genotype data were available for 1 667 children of European ethnic origin.

All parents or legal guardians gave written informed consent, and the study was approved by the IMIM-Parc de Salut Mar Research Ethics Committee (No. 2010/41221/I), Barcelona, Spain; and the FP7-ERC-2010-AdG Ethics Review Committee (268479-22022011).

*CATSS*

The Child and Adolescent Twin Study in Sweden (CATSS) is an ongoing longitudinal twin study targeting all twins born in Sweden since July 1, 1992 [4, 5]. Parents of twins are interviewed regarding the children’s somatic and mental health and social environment in connection with their 9th or 12th birthdays and followed up over time. The response rate was 75%. Follow-ups were conducted when the twins were 15 years of age (response rate, 61%) and 18 years of age (response rate, 59%). All responding parents, legal guardians or twins provided informed consent; digitally, written or by participation, to the study, and all data were deidentified. This study received ethical approval from the Karolinska Institutet Ethical Review Board. DNA samples (from saliva) were obtained from the participants at study enrollment, see Brikell *et al* [6]. A total of 11 081 samples passed quality control assessment; MZ twins were then imputed, resulting in 13 576 samples and 6 981 993 imputed SNPs that passed all quality control assessments.

*CHDS*

The New Zealand arm of the Gene-Environment-Development Initiative (GEDI) utilized data from the Christchurch Health and Development Study, a longitudinal study of the life course development of a birth cohort of 1 265 children born in the Christchurch (New Zealand) urban region in mid-1977 [7–10]. For this analysis phenotypic data on childhood aggression/ODD gathered via parental and self-report were combined with gene chip data for a sample of 626 participants of European ancestry.

*COGA*

The Collaborative Studies on the Genetics of Alcoholism (COGA) is an eleven-center research project in the United States designed to identify and understand the genetic basis of alcoholism. Research is conducted at University of Connecticut, Indiana University, University of Iowa, SUNY Downstate Medical Center at Brooklyn, Washington University in St. Louis, University of California at San Diego, Rutgers University, University of Texas Health Science Center at San Antonio, Virginia Commonwealth University, Icahn School of Medicine at Mount Sinai, and Howard University.

COGA investigators have collected data on more than 2 255 extended families in which many members are affected by alcoholism. The researchers collected extensive clinical, neuropsychological, electrophysiological, biochemical, and genetic data on the more than 17 702 individuals who are represented in the database. The researchers also have established a repository of cell lines from these individuals to serve as a permanent source of DNA for genetic studies.

*COPSAC*

The Copenhagen Prospective Studies on Asthma in Childhood is a clinical study with multiple cohorts (COPSAC2000 and COPAC2010). The COPSAC2010 cohort is a population based prospective mother-child cohort comprising 700 children born to unselected mothers (during 2009-10) from Zealland, Denmark. The cohort was enrolled at age 1 week and attended the research clinic for clinical examinations at ages 1, 3, 6, 9, 12, 18, 24, 30 and 36 month and yearly hereafter till age 8 years. The Ethics Committee for Copenhagen and the Danish Data Protection Agency approved this study. This has been described elsewhere [11]. The GWAS was run in the COPSAC 2010 birth cohort (n=459) where the phenotypes were extracted from the Strengths and Difficulty Questionnaire (SDQ) at the 8 years visit.

*Dunedin Study*

Participants were members of the Dunedin Multidisciplinary Health and Development Study, a longitudinal investigation of health and behavior in a representative birth cohort [12]. Study members (n = 1 037; 91% of eligible births; 52% male) were all individuals born between April 1972 and March 1973 in Dunedin, New Zealand, who were eligible for the longitudinal study based on residence in the province at 3 years of age and who participated in the first follow-up assessment at 3 years of age. The cohort represented the full range of socioeconomic status on NZ’s South Island. On adult health, the cohort matches the NZ National Health and Nutrition Survey (e.g., BMI, smoking, GP visits)[12]. Cohort members are primarily white; approximately 7% self-identify as having partial non-Caucasian ancestry, matching the South Island. Assessments were carried out at birth and at ages 3, 5, 7, 9, 11, 13, 15, 18, 21, 26, 32, and 38 years, when 95% of the 1 007 study members still alive took part. The Dunedin Study was approved by the NZ-HDEC (Health and Disability Ethics Committee) and informed consent was obtained from all study members.

*E-Risk*

Participants were members of E-Risk, which tracks the development of a 1994-95 birth cohort of 2,232 British children [13]. Briefly, the E-Risk sample was constructed in 1999-2000, when 1,116 families (93% of those eligible) with same-sex 5-year-old twins participated in home-visit assessments. This sample comprised 56% monozygotic (MZ) and 44% dizygotic (DZ) twin pairs; sex was evenly distributed within zygosity (49% male). The study sample represents the full range of socioeconomic conditions in Great Britain, as reflected in the families’ distribution on a neighborhood-level socioeconomic index (ACORN [A Classification of Residential Neighborhoods], developed by CACI Inc. for commercial use): 25.6% of E-Risk families live in “wealthy achiever” neighborhoods compared to 25.3% nationwide; 5.3% vs. 11.6% live in “urban prosperity” neighborhoods; 29.6% vs. 26.9% in “comfortably off” neighborhoods; 13.4% vs. 13.9% in “moderate means” neighborhoods; and 26.1% vs. 20.7% in “hard-pressed” neighborhoods. E-Risk underrepresents “urban prosperity” neighborhoods because such households are often childless.

Home visits were conducted when participants were aged 5, 7, 10, 12 and most recently, 18 years (93% participation). The Joint South London and Maudsley and the Institute of Psychiatry Research Ethics Committee approved each phase of the study. Parents gave informed written consent and twins gave written assent between 5-12 years and then informed written consent at age 18.

At age 18, 2 066 participants were assessed, each twin by a different interviewer. The average age at the time of assessment was 18.4 years (SD = 0.36); all interviews were conducted after the 18th birthday.

*FinnTwin12*

FinnTwin12 is a population-based cohort of Finnish twins born 1983–1987 that was established to track health and behavioral habits [14–16]. Identified through the Finnish Central Population Registry, families with twins were contacted and initially enrolled when the twins were 11–12 years old (N=5600 twins; 87% response rate). Questionnaires were collected from multiple informants over time: from the twins themselves at ages 12, 14, and 17; from parents at age 12; and from teachers at ages 12 and 14. DNA samples (from blood or saliva) for genotyping were taken in young adulthood (mean age 22 yrs), for which written informed consent was obtained. Study protocols were approved by the local ethics committee in Helsinki, and the IRB of Indiana University (Bloomington).

*Generation R Study*

The Generation R Study is a prospective cohort study from fetal life onwards that included pregnant women living in Rotterdam, the Netherlands, with an expected delivery date between April 2002 and January 2006 (n = 9,778). The main aim of this study is to identify early environmental and genetic factors that affect growth, health and development [17]. The Generation R Study is multidisciplinary and both prenatal and postnatal measures have included multiple domains of growth, health and development. Rotterdam is an ethnically diverse city and this is reflected in the Generation R participants. Of the enrolled mothers, 42% was of non-Dutch ethnic background, largely made by mothers from Surinamese (9%), Turkish (7%) and Moroccan (3%) background [17, 18]. Data has been collected in children up until the mean age of 10 years, with current on-going data collection at mean age 13 years. Study protocols were approved by the local ethics committee, and written informed consent and assent was obtained from all parents and children.

*GINIplus/LISA*

The influence of Life-style factors on the development of the Immune System and Allergies in East and West Germany (LISA) Study is a population-based birth cohort study. A total of 3 094 healthy, full-term neonates were recruited between 1997 and 1999 in Munich, Leipzig, Wesel and Bad Honnef. The participants were not pre-selected based on family history of allergic diseases.

A total of 5 991 mothers and their newborns were recruited into the German Infant study on the influence of Nutrition Intervention PLUS environmental and genetic influences on allergy development (GINIplus) between September 1995 and June 1998 in Munich and Wesel. Infants with at least one allergic parent and/or sibling were allocated to the interventional study arm investigating the effect of different hydrolysed formulas for allergy prevention in the first year of life. All children without a family history of allergic diseases and children whose parents did not give consent for the intervention were allocated to the non-interventional arm. Detailed descriptions of the LISA and GINIplus studies have been published elsewhere [19, 20]. DNA was collected at the age 6 and 10 years. During the 10- and 15-year follow-ups, information on aggressive behavior was collected based on the conduct problems subscale from the strength and difficulties (SDQ) questionnaire. At 10 years, the questionnaire was administered to the parents and at 15 years, to the participants themselves. For both studies, approval by the local Ethics Committees and written consent from participants and their families were obtained.

*GSMS*

The Great Smoky Mountains Study is a longitudinal, representative study of 1 420 children in 11 predominantly rural counties in Southeastern United States [21, 22]. Annual assessments on psychopathology and associated factors were completed on the 1 420 children until age 16 (6 674 observations of 1 420 individuals; 1993 to 2000) and then again at ages 19, 21, 25, and 30 (4 556 observations of 1 336 participants; 1999 to 2015) for a total of 11,230 total assessments.

*IBG*

Two cohort studies supplied information for this project: The Colorado Adoption Project (<http://ibgwww.colorado.edu/cap/> ) and the Colorado Twin Registry (<https://www.colorado.edu/ibg/research/human-research-studies/colorado-twin-registry> ). [23–25]

*INMA*

The INMA—INfancia y Medio Ambiente—(Environment and Childhood) Project is a network of birth cohorts in Spain that aim to study the role of environmental pollutants in air, water and diet during pregnancy and early childhood in relation to child growth and development (<http://www.proyectoinma.org/>) [26]. The study has been approved by Ethical Committee of each participating centre and written consent was obtained from participating parents. Data for this study comes from INMA Sabadell subcohort.

The study was approved by the Ethical Committee of the Municipal Institute of Medical Investigation and by the Ethical Committee of the hospitals involved in the study. The pregnant women received information of the study both written and orally. Their informed consent of the participants was asked in each of the visits.

*INSchool*

The INSchool cohort consist of 3, 557 children from 23 schools in Catalonia (age range: 5-17 years; mean age: 9.8 years, s.d.=2.9). 57.5% of the participants (n=2 046) were males. Screening included the Achenbach System of Empirically Based Assessment (ASEBA) with the Child Behavior Checklist CBCL/4-18 (completed by parents or surrogates), the Teacher Report Form TRF/5-18 (completed by teachers and other school staff) and the Youth Self-Report YSR/11-18 (completed by youths); the Strengths and Difficulties Questionnaire (SDQ) and the Conner’s ADHD Rating Scales (Parents and Teachers). The study was approved by the Clinical Research Ethics Committee (CREC) of Hospital Universitari Vall d'Hebron, all methods were performed in accordance to the relevant guidelines and regulations and written informed consent was obtained from participant parents before inclusion into the study.

*MCTFR*

Data for this study come from the Minnesota Center for Twin and Family Research (MCTFR), and consists of three cohorts of same-sex twins from the birth years 1971-91, 1977-85 [27], and 1988-94 [28]. Approval for all studies was obtained through the University of Minnesota Institutional Review Board.

*MOBA*

The Norwegian Mother, Father and Child Cohort Study (MoBa) is a population-based pregnancy cohort study conducted by the Norwegian Institute of Public Health [29]. Participants were recruited from all over Norway from 1999-2008. The women consented to participation in 41% of the pregnancies. The cohort now includes 114.500 children, 95.200 mothers and 75.200 fathers. The current study is based on version 11 of the quality-assured data files released for research on ADHD. The establishment of MoBa and initial data collection was based on a license from the Norwegian Data protection agency and approval from The Regional Committees for Medical and Health Research Ethics. The MoBa cohort is based on regulations based on the Norwegian Health Registry Act. The current study was approved by The Regional Committees for Medical and Health Research Ethics (number 2016/1702).

*MSUTR*

Data for this study comes from the Michigan State University Twin Registry (MSUTR), a large, population-based twin registry comprised of thousands of twins throughout Michigan, aged 3-55 (current N~31 300) [30]. The overall focus of the MSUTR is on understanding developmental changes in genetic, environmental, and neurobiological influences on internalizing and externalizing disorders. Recruitment for the MSUTR is ongoing via the identification of birth records through the Michigan Department of Health and Human Services (MDHHS). Because birth records are confidential in Michigan, recruitment packets are mailed directly from MDHHS to eligible twin pairs. Twins indicating interest in participation via pre-stamped postcards or e-mails/calls to the MSUTR project office are then contacted by study staff to determine study eligibility and to schedule their assessments. Participants in the registry complete a family health and demographic questionnaire via mail. Families are then recruited for one or more of the intensive, in-person studies based on their answers to relevant items in the registry questionnaire. In-person assessments target a variety of biological, genetic, and environmental phenotypes, including multi-informant measures of psychiatric and behavioral phenotypes, census and neighborhood informant reports of twin neighborhood characteristics, buccal swab and salivary DNA samples, assays of adolescent and adult steroid hormone levels, and/or videotaped interactions of child twin families.

Approval for all studies was obtained through the Michigan State University Institutional Review Board and the Michigan Department of Health and Human Services Institutional Review Board. All methods were performed in accordance to relevant guidelines and regulations.

*MUSP*

This study is of 7 223 women recruited early in pregnancy over the period 1981-1983 and the live singleton children to whom they subsequently gave birth. There were follow-up of the mothers at 5, 14, 21 and 27 years after recruitment. Children were also followed-up in the mothers’ questionnaire and independently at 21 and 30 years of age. The CBCL and YSR were administered to children at 5, 14 and 21 years of age. The CIDI was administered to mothers at 27 years after recruitment and the children at 21 and 30 years after recruitment. The cohort comprises both mothers (to 27 years after the birth) and children (to 30 years of age). Three papers describing the recruitment methodology and sample details have been published [31–33].

*NFBC1986*

Northern Finland Birth Cohort 1986 (NFBC1986) is a prospective longitudinal birth cohort which included pregnant women with expected date of delivery between July 1985 and June 1986 in the two Northern most provinces of Finland. In total, 9 432 children were live-born in the cohort [34]. At approximately age 16 years, the cohort members were asked to complete a postal questionnaire, including the Youth Self-Report (YSR). Items on aggression are used here. At age 16 years blood samples taken for DNA extraction for 6 266 adolescents attending the clinical examination. All participants and their parents provided consent to use their data and received Institutional Review Board approval by University of Oulu, and the Ethics committee of the Ostrobotnia Hospital district.

More information may be found at: <http://www.oulu.fi/nfbc>.

*NTR*

The Netherlands Twin Register (NTR) is a population-based prospective cohort study which includes newborn twins and multiples from the Netherlands. Recruitment started with birth year 1986 [35]. NTR data collection has a focus on growth, development, emotional and behavioral problems and health. Phenotype data on aggression were collected by surveys, in which parents and teachers were asked to rate their offspring / pupils’ behavior using standardized instruments [36, 37]. At age 14 years and after, twins and their siblings were asked for self-assessments [38]. Buccal cells and blood for DNA isolation were collected in multiple sub-projects [39]. See: <http://www.tweelingenregister.org/> for more information. The study was approved by the Central Ethics Committee on Research Involving Human Subjects of the VU University Medical Centre, Amsterdam, an Institutional Review Board certified by the U.S. Office of Human Research Protections (IRB number IRB00002991 under Federal-wide Assurance- FWA00017598; IRB/institute codes, NTR 03-180).

*QIMR*

The QIMR *Retrospective DSM III Conduct Disorder* contribution draws on data from a number of studies (phenotypic and genotypic) undertaken from the 1980s onward by the Genetic Epidemiology group at QIMR (QIMR Berghofer Medical Research Institute or QIMRB), with recruitment predominantly from families with adult twins who registered for research purposes with the Australian Twin Registry (<https://www.twins.org.au>) [40–42]. Phenotype data were self-report, with study overlap resolved by using the questionnaire completed at the youngest age. The largest contributions were from (1) SS1, an interview study of adult twins born before 1964 using the Semi-Structured Assessment for the Genetics of Alcoholism (SSAGA) instrument (N=4 046 twins used, conducted 1993-1995) and SP, a follow-up of their spouses (N=584, conducted 1998-1999); (2) Twin-89, an interview study on personality and drinking habits of the twins born 1964-1972 (N=2 040, conducted 1996-2000); (3) the NIH-funded Nicotine Addiction Genetics (NAG) and three Interactive Research Project Grant (IRPG) interview studies (N=4 017, from both cohorts, conducted 2003-2005). Closely similar or identical questions approximating the items in the DSM-III CD diagnosis (testing aggressive and highly anti-social behavior), were scored for all studies and combined into a 14 or 15-item symptom score, rescaled by 15/14 if there were 14 items.

TCHAD

The twin study of Child and Adolescent Development (TCHAD) followed 1 500 twin pairs from age 8 to age 26 [5]. The genotyping followed the same procedure as for CATSS; see Brikell *et al* [6].

*The Raine Study*

The Raine Study is a prospective pregnancy cohort where 2 900 mothers (Gen1) where recruited between 1989 and 1991 [43, 44]. Recruitment took place at Western Australia’s major perinatal centre, King Edward Memorial Hospital, and nearby private practices. Women who had sufficient English language skills, an expectation to deliver at King Edward Memorial Hospital, and an intention to reside in Western Australia to allow for future follow-up of their child (Gen2) were eligible for the study.

The Raine Study is known to be one of the largest successfully prospective cohorts richly phenotyped at multiple time points over pregnancy, infancy, childhood adolescence, and young adult. The mothers (Gen1) completed questionnaires regarding their children (Gen2) and the children (Gen2) had physical examinations at ages 1, 2, 3, 6, 8, 10, 14, 17, 20 and 22 years.

*TEDS*

The Twins Early Development Study (TEDS) is a longitudinal twin study that recruited over 16 000 twin pairs born between 1994 and 1996 in England and Wales through national birth records [45]. More than 10 000 of these families are still involved in study. TEDS was and still is a representative sample of the population in England and Wales. Rich cognitive and behavioural data have been collected from the twins from infancy to emerging adulthood with data collection at ages 2, 3, 4, 7, 8, 9, 10, 12, 14, 16, 18, 19 and 21, enabling longitudinal genetically sensitive study designs. Data have been collected from twins themselves (including extensive web-based cognitive testing), from their parents and teachers, and from the UK National Pupil Database. Genotyped DNA data are available for 10 346 individuals (who are unrelated except for 3 320 dizygotic co-twins). TEDS data have contributed to over 400 scientific papers involving more than 140 researchers in 50 research institutions.

*TRAILS*

Tracking Adolescents’ Individual Lives Survey (TRAILS) is a large prospective population study of Dutch adolescents with bi- or triennial measurements from age 11 years onwards. TRAILS participants were selected from five municipalities in the Northern part of the Netherlands [46]. The cohort’s characteristics and database are described in detail elsewhere [47] and at <http://www/trails.nl/>. DNA was extracted from blood samples or (in a few cases) buccal swaps, collected at about age 16. In total,

1 491 children of white European descent were genotyped. The study was approved by the Dutch Central Committee on Research Involving Human subjects (CCMO), and all measurements were carried out with participants’ adequate understanding and written consent.

*VTSABD*

The VCU arm of the NIDA-funded Gene-Environment-Development Initiative (GEDI) combined existing phenotypic and environmental data from the Virginia Twin Study of Adolescent Behavioral Development (VTSABD) study, a population-based multi-wave, cohort-sequential twin study of adolescent psychopathology and its risk factors, with genome-wide genotyping, generating a genotyped sample of ~900 subjects [10, 48–51].

*YFS*

The Young Finns study (YFS) is an on-going longitudinal population-based cohort study that includes

3 596 healthy Finnish children and adolescents from six birth cohorts (aged 3, 6, 9, 12, 15, and 18 years at the study baseline in 1980)[52, 53]. The study was approved by the ethical committee of the Varsinais-Suomi’s hospital district’s federation of municipalities. For the current study, we selected a subsample of 1 784 participants from the five oldest age groups (aged 6, 9, 12, 15, and 18 at baseline) who had parent-reported aggressive behavior data and genetic data available.

**Acknowledgment and Funding**

This work is supported by the "Aggression in Children: Unraveling gene-environment interplay to inform Treatment and InterventiON strategies" (ACTION) project. ACTION receives funding from the European Union Seventh Framework Program (FP7/2007-2013) under grant agreement no 602768.

*ABCD*

We thank all participating hospitals, obstetric clinics, general practitioners and primary schools for their assistance in implementing the ABCD study. We also gratefully acknowledge all the women and children who participated in this study for their cooperation.

The ABCD study has been supported by grants from The Netherlands Organisation for Health Research and Development (ZonMW) and Sarphati Amsterdam. Genotyping was funded by the BBMRI-NL grant CP2013-50. Dr M.H. Zafarmand was supported by BBMRI-NL (CP2013-50). Dr. T.G.M. Vrijkotte was supported by ZonMW (TOP 40–00812–98–11010).

*ALSPAC*

We are extremely grateful to all the families who took part in this study, the midwives for their help in recruiting them, and the whole ALSPAC team, which includes interviewers, computer and laboratory technicians, clerical workers, research scientists, volunteers, managers, receptionists and nurses. GWAS data were generated by Sample Logistics and Genotyping Facilities at Wellcome Sanger Institute and LabCorp (Laboratory Corporation of America) using support from 23andMe.

The UK Medical Research Council and Wellcome (Grant ref: 102215/2/13/2) and the University of Bristol provide core support for ALSPAC. A comprehensive list of grants funding for ALSPAC is available online (http://www.bristol.ac.uk/alspac/external/documents/grant-acknowledgements.pdf).

GDS works in a unit that receives funding from the University of Bristol and the UK Medical Research Council (MC_UU_00011/1). BSTP is supported through Max Planck Society core funding and the Simons Foundation (514787).

*BREATHE*

We acknowledge all the families and schools participating in the study. The research leading to these results has received funding from the European Research Council under the ERC Grant Agreement number 268479 – the BREATHE project. ISGlobal is a member of the CERCA Programme, Generalitat de Catalunya. We thank the La Caixa Foundation for their financial support in the PAHs analyses. S. Alemany is funded by avJuan de la Cierva – Incorporación Postdoctoral Contract from Ministerio de Economía, Industria y Competitividad (IJCI-2017-34068).

*CATSS*

The Child and Adolescent Twin Study in Sweden study was supported by the Swedish Council for Working Life, funds under the ALF agreement, the Söderström Königska Foundation and the Swedish Research Council (Medicine, Humanities and Social Sciences, and SIMSAM). We acknowledge The Swedish Twin Registry for access to data which is managed by Karolinska Institutet and receives funding through the Swedish Research Council under the grant no 2017-00641.

*CHDS (GEDI)*

The Christchurch Health and Development Study has been supported by funding from the Health Research Council of New Zealand, the National Child Health Research Foundation (Cure Kids), the Canterbury Medical Research Foundation, the New Zealand Lottery Grants Board, the University of Otago, the Carney Centre for Pharmacogenomics, the James Hume Bequest Fund, US National Institutes of Health grant MH077874 and National Institute on Drug Abuse grant R01DA024413.

*COGA*

The Collaborative Study on the Genetics of Alcoholism (COGA), Principal Investigators B. Porjesz, V. Hesselbrock, H. Edenberg, L. Bierut, includes eleven different centers: University of Connecticut (V. Hesselbrock); Indiana University (H.J. Edenberg, J. Nurnberger Jr., T. Foroud); University of Iowa (S. Kuperman, J. Kramer); SUNY Downstate (B. Porjesz); Washington University in St. Louis (L. Bierut, J. Rice, K. Bucholz, A. Agrawal); University of California at San Diego (M. Schuckit); Rutgers University (J. Tischfield, A. Brooks); Department of Biomedical and Health Informatics, The Children’s Hospital of Philadelphia; Department of Genetics, Perelman School of Medicine, University of Pennsylvania, Philadelphia PA (L. Almasy), Virginia Commonwealth University (D. Dick), Icahn School of Medicine at Mount Sinai (A. Goate), and Howard University (R. Taylor). Other COGA collaborators include: L. Bauer (University of Connecticut); J. McClintick, L. Wetherill, X. Xuei, Y. Liu, D. Lai, S. O’Connor, M. Plawecki, S. Lourens (Indiana University); G. Chan (University of Iowa; University of Connecticut); J. Meyers, D. Chorlian, C. Kamarajan, A. Pandey, J. Zhang (SUNY Downstate); J.-C. Wang, M. Kapoor, S. Bertelsen (Icahn School of Medicine at Mount Sinai); A. Anokhin, V. McCutcheon, S. Saccone (Washington University); J. Salvatore, F. Aliev, B. Cho (Virginia Commonwealth University); and Mark Kos (University of Texas Rio Grande Valley). A. Parsian and M. Reilly are the NIAAA Staff Collaborators.

We continue to be inspired by our memories of Henri Begleiter and Theodore Reich, founding PI and Co-PI of COGA, and also owe a debt of gratitude to other past organizers of COGA, including Ting-Kai Li, P. Michael Conneally, Raymond Crowe, and Wendy Reich, for their critical contributions. This national collaborative study is supported by NIH Grant U10AA008401 from the National Institute on Alcohol Abuse and Alcoholism (NIAAA) and the National Institute on Drug Abuse (NIDA), the NIH K02 Award to Dr. Danielle Dick K02 AA018755 from the National Institute on Alcohol Abuse and Alcoholism (NIAAA).

*COPSAC*

All funding received by COPSAC is listed on www.copsac.com. The Lundbeck Foundation (Grant no R16-A1694); The Ministry of Health (Grant no 903516); Danish Council for Strategic Research (Grant no 0603-00280B) and The Capital Region Research Foundation have provided core support to the COPSAC research center. We express our deepest gratitude to the children and families of the COPSAC 2010 cohort study for all their support and commitment. We acknowledge and appreciate the unique efforts of the COPSAC research team.

*Dunedin*

We thank the Dunedin Study members and their parents, Unit research staff, and Study founder Phil Silva. The Dunedin Longitudinal Study is funded by the New Zealand Health Research Council, the New Zealand Ministry of Business, Innovation, and Employment (MBIE), the National Institute on Aging (AG032282), and the Medical Research Council (MR/P005918/1). Additional support was provided by the Jacobs Foundation and the Avielle Foundation. This work used a high-performance computing facility partially supported by grant 2016-IDG-1013 (“HARDAC+: Reproducible HPC for Next-generation Genomics”) from the North Carolina Biotechnology Center.

*E-Risk*

We are grateful to the study mothers and twins for their participation, and to members of the E-Risk team for their dedication, hard work, and insights. The E-Risk Study is funded by the Medical Research Council (G1002190) and the National Institute of Child Health and Human Development (HD077482). Additional support was provided by the Jacobs Foundation. This work used a high-performance computing facility partially supported by grant 2016-IDG-1013 (“HARDAC+: Reproducible HPC for Next-generation Genomics”) from the North Carolina Biotechnology Center.

LA is appointed Mental Health Leadership Fellow for the UK Economic and Social Research Council (ESRC)

*FinnTwin12*

Data collection has been supported by the National Institute of Alcohol Abuse and Alcoholism (Grants AA-12502, AA-00145, and AA-09203 to RJR) and the Academy of Finland (Grants 100499, 205585, 118555, 141054, 265240, 263278 and 264146 to JK). Genotyping has been supported by grants AA15416 and K02AA018755 to D M Dick, and Academy of Finland Center of Excellence in Complex Disease Genetics (grant numbers: 213506, 129680 to JK). JK has been supported by the Academy of Finland (Grant 312073). AW is supported by the "Aggression in Children: Unraveling gene-environment interplay to inform Treatment and InterventiON strategies" (ACTION) project. ACTION receives funding from the European Union Seventh Framework Program (FP7/2007-2013) under grant agreement no 602768.

We wish to sincerely thank all of the twins and their families, school principals, and teachers who participated in the FinnTwin12 study, and the FinnTwin12 data collection staff for all their hard work.

*Generation R Study*

This work was supported by the Dutch Ministry of Education, Culture and Science (Gravity Grant No. 024.001.003, Consortium on Individual Development), and the Netherlands Organisation for Scientific Research (NWO‐grant 016.VICI.170.200) to HT. The first phase of the Generation R Study is made possible by financial support from the Erasmus Medical Centre, Rotterdam; the Erasmus University Rotterdam; and the Netherlands Organisation for Health Research and Development (ZonMw). The authors gratefully acknowledge the contribution of all children and parents, general practitioners, hospitals, midwives and pharmacies involved in the Generation R Study. The Generation R Study is conducted by the Erasmus Medical Centre (Rotterdam) in close collaboration with the School of Law and Faculty of Social Sciences of the Erasmus University Rotterdam; the Municipal Health Service Rotterdam area, Rotterdam; the Rotterdam Homecare Foundation, Rotterdam; and the Stichting Trombosedienst & Artsenlaboratorium Rijnmond, Rotterdam.

*GINIplus/LISA*

The authors thank all families for participation in the studies and the LISA and GINIplus study teams for their excellent work.

*GSMS (GEDI)*

This research was supported by the National Institute on Drug Abuse (U01DA024413, R01DA11301), the National Institute of Mental Health (R01MH063970, R01MH063671, R01MH048085, K01MH093731 and K23MH080230), NARSAD, and the William T. Grant Foundation. We are grateful to all the GSMS and CCC study participants who contributed to this work.

*IBG*

Funding for genotyping, analytic support, and data curation supported by NIH R01 AG046938 and P60 DA011015. Dr. Hopfer reports support from DA032555, DA035804, and DA042755.

*INMA*

This work was supported by grants from the European Union [FP7-ENV-2011 cod 282957 and HEALTH.2010.2.4.5-1] and from Spain: Instituto de Salud Carlos III [Red INMA G03/176, CB06/02/0041, FIS-FEDER: PI03/1615, PI041436, PI04/1509, PI04/1112, PI04/1931, PI05/1079, PI05/1052, PI06/0867, PI06/1213, PI07/0314, PI081151, PI09/02647, PI09/00090, PI11/01007, PI11/02591, PI11/02038, PI12/01890, PI13/1944, PI13/2032, PI14/00891, PI14/01687, PI16/1288, and PI17/00663; Miguel Servet-FEDER CP11/0178CP15/0025; Miguel Servet-FSE MS16/00128, MS15/0025, and MSII16/00051; and PFIS-FI14/00099], Alicia Koplowitz Foundation 2017, Generalitat Valenciana, , Department of Health of the Basque Government [2005111093 and 2009111069], the Provincial Government of Gipuzkoa [DFG06/004 and DFG08/001], and the Generalitat de Catalunya-CIRIT [1999SGR 00241].

The authors are grateful to the mothers and children who participated in the study.

*INSchool*

We are grateful to all the families and schools who kindly participated in the study. This work was funded by the Instituto de Salud Carlos III (PI16/01505, PI17/00289, PI18/01788, PI19/01224 and PI19/00721), and co‐financed by the European Regional Development Fund (ERDF), Agència de Gestió d’Ajuts Universitaris i de Recerca- AGAUR, Generalitat de Catalunya, Spain (2014SGR1357, 2017SGR1461), the Health Research and Innovation Strategy Plan (PERIS SLT006/17/285 and PERIS SLT006/17/287), Generalitat de Catalunya, Spain, la Fundació Bancària ”La Caixa”, els Departaments de Salut i d’Educació, Generalitat de Catalunya, Spain, les Diputacions de Barcelona i Lleida, Spain, the European College of Neuropsychopharmacology (ECNP network: ‘ADHD across the lifespan’) and a NARSAD Young Investigator Grant from the Brain & Behavior Research Foundation. The research leading to these results has received funding from the European Union Seventh Framework Program (FP72007-2013) under grant agreement No 602805 and from the European Union H2020 Programme (H2020/2014-20) under grant agreements Nos. 667302 (CoCA) and 728018 (Eat2BeNICE). Over the course of this investigation, M. Ribases was a recipient of a Miguel de Servet contract from the Instituto de Salud Carlos III, Spain (CP09/00119 and CPII15/ 00023), P. Rovira was a recipient of a pre-doctoral fellowship from the Agència de Gestió d’Ajuts Universitaris i de Recerca (AGAUR), Generalitat de Catalunya, Spain (2016FI_B 00899), C. Sánchez-Mora was a recipient of a Sara Borrell contract and a mobility grant from the Spanish Ministerio de Economía y Competitividad, Instituto de Salud CarlosIII (CD15/00199 and MV16/00039) and M. Soler Artigas was a recipient of a contract from the Biomedical Network Research Center on Mental Health (CIBERSAM), Madrid, Spain.

M.C. has received travel grants and research support from Eli Lilly and Co., Janssen-Cilag, Shire and Lundbeck and served as consultant for Eli Lilly and Co., Janssen-Cilag, Shire and Lundbeck.

J.A.R.Q was on the speakers’ bureau and/or acted as consultant for Eli-Lilly, Janssen-Cilag, Novartis, Shire, Lundbeck, Almirall, Braingaze, Sincrolab, Medice, Exeltis and Rubió in the last 5 years. He also received travel awards (air tickets + hotel) for taking part in psychiatric meetings from Janssen-Cilag, Rubió, Shire, Medice and Eli- Lilly. The Department of Psychiatry chaired by him received unrestricted educational and research support from the following companies in the last 5 years: Eli-Lilly, Lundbeck, Janssen- Cilag, Actelion, Shire, Ferrer, Oryzon, Roche, Psious, and Rubió.

*MCTFR*

MCTFR research is supported by grants from the National Institute on Drug Abuse (R37 DA005147, R01 DA013240, R01 DA036216, and U01 DA024417), the National Institute on Alcohol Abuse and Alcoholism (R37 AA009367 and R01 AA011886), and the National Institute of Mental Health (R01 MH066140).

*MoBa*

The Norwegian Mother, Father and Child Cohort Study is supported by the Norwegian Ministry of Health and Care Services and the Ministry of Education and Research. We are grateful to all the participating families in Norway who take part in this on-going cohort study. We thank the Norwegian Institute of Public Health (NIPH) for generating high-quality genomic data. This research is part of the HARVEST collaboration, supported by the Research Council of Norway (#229624). We also thank the NORMENT Centre for providing genotype data, funded by the Research Council of Norway (#223273), South East Norway Health Authority and KG Jebsen Stiftelsen. We further thank the Center for Diabetes Research, the University of Bergen for providing genotype data funded by the ERC AdG project SELECTionPREDISPOSED, Stiftelsen Kristian Gerhard Jebsen, Trond Mohn Foundation, the Research Council of Norway, the Novo Nordisk Foundation, the University of Bergen, and the Western Norway health Authorities (Helse Vest).

The analyses of the data were supported by Stiftelsen Kristian Gerhard Jebsen (grant number SKGJ‐MED‐ 002) and by NIH/NIMH (grant number 5U01MH109539‐03).

*MSUTR*

This research was supported by the National Institute of Mental Health (NIMH) under Award Number R01-MH081813 and the Eunice Kennedy Shriver National Institute for Child Health and Human Development (NICHD) under Award Number R01-HD066040

This research was supported by the National Institute on Drug Abuse under Award Number R01DA043501 and the National Library of Medicine under Award Number R01LM012848

*MUSP*

This study was funded by grants received from the National Health and Medical Research Council (NHMRC) and Australian Research Council (ARC). Thanks also to Shelby Marrington (Project Manager) and Greg Shuttlewood (Data Manager) who have supervised the day-to-day management of the study. We also extend our thanks to the mothers and children who have continued to participate in the study.

JGS is supported by a National Health and Medical Research Council Practitioner Fellowship Grant

APP1105807

The authors thank the MUSP study participants and study team. The authors thank the National Health and Medical Research Council (NHMRC).

*NFBC*

We thank all cohort members and researchers who have participated in the study. We also wish to acknowledge the work of the NFBC project center. NFBC1986 has received funding from: EU QLG1-CT-2000-01643 (EUROBLCS) Grant no. E51560, NorFA Grant no. 731, 20056, 30167, USA / NIHH 2000 G DF682 Grant no. 50945, the EU H2020-MSCA-ITN-2016 CAPICE Action Grant no. 721567. Academy of Finland EGEA project (285547), EU H2020 LifeCycle Action (grant agreement No 733206), and DynaHEALTH action (grant agreements No. 633595). The DNA extractions, sample quality controls, biobank upkeep and aliquoting were performed in the National Public Health Institute, Biomedicum Helsinki, Finland and supported financially by the Academy of Finland and Biocentrum Helsinki.

*NTR*

Funding was obtained from multiple grants from the Netherlands Organization for Scientific Research (NWO) and The Netherlands Organisation for Health Research and Development (ZonMW): Genetic influences on stability and change in psychopathology from childhood to young adulthood (ZonMw 912-10-020); Twin family database for behavior genomics studies (NWO 480-04-004); Genetic and Family Influences on Adolescent. Psychopathology and Wellness (NWO 463-06-001); A Twin-Sibling Study of Adolescent Wellness (451-04-034)**.** Twin research focusing on behavior (NWO 400-05-717); Longitudinal data collection from teachers of Dutch twins and their siblings (481-08-011), Twin-family-study of individual differences in school achievement (NWO-FES, 056-32-010), Genotype/phenotype database for behavior genetic and genetic epidemiological studies (ZonMw Middelgroot 911-09-032); “Why some children thrive” (OCW_Gravity program –NWO-024.001.003), Netherlands Twin Registry Repository: researching the interplay between genome and environment (NWO-Groot 480-15-001/674); BBMRI –NL (184.021.007 and 184.033.111): Biobanking and Biomolecular Resources Research Infrastructure; Spinozapremie (NWO- 56-464-14192) and KNAW Academy Professor Award (PAH/6635) to DIB ; the Neuroscience Campus Amsterdam (NCA) and Amsterdam Public Health (APH); the European Science Council (ERC) Genetics of Mental Illness (ERC Advanced, 230374); NIH: Rutgers University Cell and DNA Repository cooperative agreement (NIMH U24 MH068457-06); Developmental Study of Attention Problems in Young Twins (NIMH, RO1 MH58799-03); Grand Opportunity grant Developmental trajectories of psychopathology (NIMH 1RC2 MH089995) and the Avera Institute for Human Genetics.

*QIMR*

Phenotype and genotype collection was funded by the National Health and Medical Research Council (including grants APP1103603, 241944, 339462, 389927,389875, 389891, 389892, 389938, 442915, 442981, 496739, 552485,and 552498); by the Australian Research Council (including grants A7960034, A79906588, A79801419, DP0770096, DP0212016, and DP0343921), and National Institutes of Health (including grants AA013320, AA013321, AA013326, AA011998 and AA017688). The authors acknowledge the extensive work carried out by current and former QIMRB staff, particularly the current QIMRB Sample Processing facility (formerly Molecular Epidemiology lab) for sample processing; former interviewers, IT and project staff for recruitment and data collection; and the use of the QIMRB High Performance Computing facility for data storage and analysis. SEM is supported by an NHMRC Senior Research Fellowship (APP1103623) LC-C is supported by a QIMR Berghofer Fellowship.

*The Raine Study*

The Raine Study acknowledges the National Health and Medical Research Council (NHMRC) for their long term contribution to funding the study over the last 29 years. Core Management of the Raine Study has been funded by the University of Western Australia (UWA), Curtin University, the UWA Faculty of Medicine, Dentistry and Health Sciences, the Raine Medical Research Foundation, the Telethon Kids Institute, the Women and Infants Research Foundation, Edith Cowan University, Murdoch University, and the University of Notre Dame. This study was supported by the National Health and Medical Research Council of Australia [grant numbers 572613, 403981 and 003209] and the Canadian Institutes of Health Research [grant number MOP-82893]. The authors gratefully acknowledge the assistance of the Western Australian DNA Bank (National Health and Medical Research Council of Australia National Enabling Facility). All analytic work was supported by resources provided by the Pawsey Supercomputing Centre with funding from the Australian Government and the Government of Western Australia.

*TCHAD*

The Swedish Twin study of Child and Adolescent Development (TCHAD) was supported by the Swedish Council for Working Life and the Swedish Research Council (Medicine and SIMSAM).

*TEDS*

We gratefully acknowledge the ongoing contribution of the participants in the Twins Early Development Study (TEDS) and their families. TEDS is supported by a program grant to RP from the UK Medical Research Council (MR/M021475/1 and previously G0901245), with additional support from the US National Institutes of Health (AG046938). RP is supported by a Medical Research Council Professorship award (G19/2).

*TRAILS*

TRAILS (TRacking Adolescents’ Individual Lives Survey) is a collaborative project involving various departments of the University Medical Center and University of Groningen, the University of Utrecht, the Radboud Medical Center Nijmegen, and the Parnassia Bavo group, all in the Netherlands. TRAILS has been financially supported by grants from the Netherlands Organization for Scientific Research NWO (Medical Research Council program grant GB-MW 940-38-011; ZonMW Brainpower grant 100-001-004; ZonMw Risk Behavior and Dependence grant 60-60600-97-118; ZonMw Culture and Health grant 261-98-710; Social Sciences Council medium-sized investment grants GB-MaGW 480-01-006 and GB-MaGW 480-07-001; Social Sciences Council project grants GB-MaGW 452-04-314 and GB-MaGW 452-06-004; NWO large-sized investment grant 175.010.2003.005; NWO Longitudinal Survey and Panel Funding 481-08-013 abd 481-11-001; NWO Vici 016.130.002 and 453-16-007/2735; NWO Gravitation 024.001.003); the Dutch Ministry of Justice (WODC), the European Science Foundation (EuroSTRESS project FP-006), the European Research Council (ERC-2017-STG-757364 en ERC-CoG-2015-681466), Biobanking and Biomolecular Resources Research Infrastructure BBMRI-NL (CP 32), the Gratama foundation, the Jan Dekker foundation, the participating universities, and Accare Center for Child and Adolescent Psychiatry.

We are grateful to all adolescents, their parents and teachers who participated in this research and to everyone who worked on this project and made it possible. Statistical analyses were carried out on the Genetic Cluster Computer (http://www.geneticcluster.org), which is financially supported by the Netherlands Scientific Organization (NWO 480-05-003) along with a supplement from the Dutch Brain Foundation.

*VTSABD (GEDI)*

This research was supported by the National Institute on Drug Abuse (U01DA024413, R01DA025109), the National Institute of Mental Health (R01MH045268, R01MH068521). We are grateful to all the VTSABD study participants who contributed to this work.

*YFS*

The Young Finns Study has been financially supported by the Academy of Finland: grants 322098, 286284, 134309 (Eye), 126925, 121584, 124282, 129378 (Salve), 117787 (Gendi), and 41071 (Skidi); the Social Insurance Institution of Finland; Competitive State Research Financing of the Expert Responsibility area of Kuopio, Tampere and Turku University Hospitals (grant X51001); Juho Vainio Foundation; Paavo Nurmi Foundation; Finnish Foundation for Cardiovascular Research ; Finnish Cultural Foundation; The Sigrid Juselius Foundation; Tampere Tuberculosis Foundation; Emil Aaltonen Foundation; Yrjö Jahnsson Foundation; Signe and Ane Gyllenberg Foundation; Diabetes Research Foundation of Finnish Diabetes Association; EU Horizon 2020 (grant 755320 for TAXINOMISIS); European Research Council (grant 742927 for MULTIEPIGEN project); and Tampere University Hospital Supporting Foundation.

We thank the teams that collected data at all measurement time points; the persons who participated as both children and adults in these longitudinal studies; and biostatisticians Irina Lisinen, Johanna Ikonen, Noora Kartiosuo, Ville Aalto, and Jarno Kankaanranta for data management and statistical advice.

**Supplementary Methods**

**Sample description**

Cohorts with assessment of AGG in children and adolescents aged 1.5 - 18 years – and adult retrospective assessment of adolescent CD – with genotyping, that collaborate within the ACTION (Aggression in Children: unraveling gene-environment interplay to inform Treatment and InterventiON strategies) ( 54, 55) and the EAGLE (EArly Genetics and Lifecourse Epidemiology; Middeldorp et al. 2019) consortia and additional childhood cohorts took part in the meta-analysis. Each cohort received a standard operation protocol ( <https://www.action-euproject.eu/content/data-protocols>). Cohorts could contribute one or multiple GWASs for AGG, for every combination of rater, instrument, and age. For more information on the cohorts see Supplementary Table 1 and Supplementary Text. As limited data was available on individuals of non-European ancestry and to avoid inducing population stratification, we restricted the analysis to individuals of European ancestry. In total, 29 cohorts contributed 164 GWASs, resulting in a total of 328 935 observations on 87 485 unique individuals (Supplementary Table 2).

**Measurement of Aggressive Behavior**

AGG was rated by mothers, fathers, teachers, and by self-report. GWASs where no distinction was made between mother or father report were included with mother-reported data. In total, the meta-analysis included AGG assessed with 26 instruments (see Supplementary Table 3). The instruments covered various childhood behaviors and disorders with an aggressive component (e.g. conduct disorder and oppositional defiant disorder). Items included content such as “hot tempers” and “gets in many fights”. For all instruments and raters, AGG was assessed on a continuous scale, with higher scores indicating higher levels of AGG. The most commonly employed instruments came from the Achenbach System of Empirically Based Assessment (ASEBA; 41%; Achenbach et al. 2017) and the Strengths and Difficulties Questionnaire (SDQ; 34%; Goodman 2001).

**Genotyping and quality control**

Genotyping was performed within each cohort on common genotyping arrays (see Supplementary Table 4), followed by cohort-specific quality control based on variant- and individual-based call rate, minor allele frequency, Hardy-Weinberg equilibrium, and excessive heterozygosity (see Supplementary Table 5). Cohorts removed samples with non-European ancestry or mismatched sex. The most commonly used genotyping array across cohorts were the Illumina 660K, Illumina 670K, and Affymetrix 6.0 arrays. Cohorts were asked to use genotypes imputed to the 1000 Genomes (1000G) reference set, mapped to build 37 of the human Genome Reference Consortium assembly (GRCh37). 75.9% of the cohorts used 1000G phase 3 version 5 as reference set for the imputation, while the remaining ones imputed to 1000G phase 1 version 3 (see Supplementary Table 6).

**GWAS model**

For all cohorts, local analysts performed univariate GWASs where AGG was regressed on the SNP with sex, age, and first five ancestry-based principal components as covariates, and, if necessary, cohort-specific covariates (see Supplementary Table 7). GWASs included autosomal SNPs. Cohorts with a sample that included only unrelated subjects applied a linear regression model. To correct for non-independence of observations, cohorts with a sample containing related individuals applied a mixed linear model. Alternatively, cohorts with a sample containing related individuals could apply a sandwich correction of the standard errors [59].

Genome-wide association analyses were stratified by (1) rater, (2) instrument, and (3) age, selecting strata such that every stratum contained at least 450 observations. In total, summary statistics for 164 GWASs analyses were uploaded. Descriptive statistics for each uploaded GWAS are shown in Supplementary Table 8. Each cohort supplied the phenotypic correlations (Supplementary Table 10) between the AGG measures within the different strata and the degree of sample overlap between strata (Supplementary Table 11). These statistics allowed us to account for dependence within cohort (see “Meta-analysis method”).

**Pre-GWAMA QC**

Each uploaded file underwent QC using the EasyQC software package [60]. SNPs with a genotyping rate below 95% were removed. Additionally, based on the sample size of the GWAS, we applied variable QC filter on MAF and HWE *p*-value (see Supplementary Figure 1). A high-pass cutoff of 0.6 and 0.7 was applied to SNPs that were imputed using MACH and IMPUTE, respectively [61]. Reported allele frequencies were compared with an imputation-matched reference file (see URLs) and variants with an absolute difference larger than 0.2 were removed. Supplementary Table 9 reports the number of remaining SNPs before and after QC.

**Meta-analysis method**

Within cohort, the measures of AGG may be dependent as a result of including repeated measures and/or AGG as assessed by multiple raters. Any covariance between test statistics within cohort and across GWASs is therefore a sum of (1) a truly shared genetic signal and (2) sample overlap [62]:

$$E\left[ Z_{ji}Z_{jk}|\mathcal{l}_{j} \right]=\frac{\sqrt{N_{ji}N_{jk}}\varrho_{\mathcal{g}}}{M}\mathcal{l}_{j}+\frac{N_{s}r_{p}}{\sqrt{N_{ji}N_{jk}}}$$

Note, this function is a sum of two elements: the truly shared genetic effect on the left side of the plus sign and the cross-trait-intercept (CTI) on the right side. The CTI reflects the expected dependence between two GWASs that arises from sample overlap and phenotypic correlation. Here $Z_{ji}$ and $Z_{jk}$ are the $Z$-scores for SNP $j$ in GWASs $i$ and $k$, respectively; $\mathcal{l}_{j}=\sum_{l} r_{jl}^{2}$ is the so-called “LD Score” of SNP $j$, which measures the amount of genetic variation tagged by $j$; $N_{ji}$ and $N_{jk}$ represent the sample sizes; $\varrho_{g}$ is the genetic covariance between the traits analyzed in $i$ and $k$; $M$ stands for the number of overlapping SNPs; $N_{s}$ indicates the number of individuals that participated in both $i$ and $k$; and $r_{p}$ is the phenotypic correlation between $i$ and $k$. The CTI between $i$ and $k$ is then:

$${CTI}_{ik}=\frac{N_{s}r_{p}}{\sqrt{N_{ji}N_{jk}}}$$

To account for the effect of sample overlap, we applied a modified version of the multivariate meta-analysis approach developed by Baselmans *et al* (2019): instead of estimating the CTI using Linkage Disequilibrium SCore regression (LDSC; Bulik-Sullivan et al. 2015b, a), we calculated the expected CTI using the observed sample overlap and phenotypic covariance as reported by the cohorts. The multivariate test statistics are obtained from:

$$Z_{multi,j}=\frac{\sum_{i=1}^{P} w_{ji}Z_{ji}}{\sqrt{\sum_{i=1}^{P} w_{ji}V_{ji}+\sum_{i=1}^{P} \sum_{k=1}^{P} \sqrt{w_{ji}w_{jk}}{CTI}_{ik} for i\neq k}}$$

where $P$ is the number of GWASs across which we run the meta-analysis; $w_{ji}=\sqrt{N_{ji}h_{SNP,i}^{2}}$ is the weight given to the $j$th SNP in GWAS $i$, with $h_{SNP,i}^{2}$ being the SNP-heritability of the trait analyzed in GWAS $i$; and $V_{ji}=1$ represents the variance of the distribution of $Z_{ji}$ under the null hypothesis of no effect. Finally, we approximate the effective sample size (N_eff_) via:

$$N_{eff}={\sqrt{N}^{T}CTI}^{-1}\sqrt{N}$$

where $N$ is an $P$-sized vector of sample sizes, and $CTI$ is the $P$ x $P$ matrix of cross-trait-intercepts. When there is no sample overlap (or a phenotypic correlation equal to zero) between the GWASs (i.e. $CTI$ is an identity matrix), N_eff_ is equal to the sum of sample sizes.

**Creating age-bins for the age-specific GWAMAs**

Age-bins for the age-by-rater GWAMAs were created such that the total *univariate* number of observations exceeded 15 000. To do so, the GWAs were sorted on their mean age, from lowest to highest. For the first age-bin, the first GWA was used as starting point. Then, if the next GWA belonged to another cohort, this was included in the age-bin. If the next GWA belonged to the same cohort, the GWA with the largest sample size was retained in the age-bin. This was repeated until the total univariate N_obs_ exceeded 15 000. Then, the lower bound of the age-bin was set to the mean age of the first GWA, rounded down to the nearest integer and the upper bound of the age-bin was set to the mean age of the last GWA, rounded up to the nearest integer. Finally, all GWAs with a mean age within the boundaries (up to and including the upper bound) were included in the age-by-rater GWAMA. For the next age-bin, the first available GWA was used as the starting point and the process was repeated until all GWAs were divided into age-bins. If the last age-bin contained N_obs_ < 15 000, the last two age-bins were combined.

**Calculating polygenic scores**

All data were meta-analyzed twice more, once omitting all data from the Netherlands Twin Register (NTR) and once omitting all Australian data (Queensland Institute for Medical Research [QIMR] and Mater-University of Queensland Study of Pregnancy [MUSP]). For the target sample in the NTR we considered mother-reported AGG at age 7 (*N*=4,491), which represents the largest NTR univariate stratum. In the QIMR participants (*N*=10,706), we tested whether our childhood AGG polygenic scores (PGS) predicted adult retrospective assessment of their own CD behavior during adolescence. We allowed for cohort-specific best practice in the polygenic score analysis.

In the NTR, we created 16 sets of PGSs in PLINK1.9 [65], with *P*-value thresholds between 1 and 1.0E-05 (Supplementary Table 13). The remaining SNPs were clumped in PLINK. We applied an $r^{2}$-threshold (high-pass) of 0.5 and minimum clumping distance of 250,000 base pair positions [65]. Age; age^2^; sex; first five ancestry-based principal components (PCs); a SNP-array variable; and interaction terms between sex and age, and sex and age^2^ were defined as fixed effects. To account for relatedness, prediction was performed using generalized equation estimation as implemented in the “gee” package (version 4.13-19) in R (version 3.5.3)[66]. GEE applies a sandwich correction over the standard errors to account for clustering in the data [67]. To correct for multiple testing, we applied an FDR correction at $\alpha$=0.05 for 16 tests.

QIMR excluded SNPs with low imputation quality ($r^{2}$=0.6) and MAF below 1%, and selected the most significant independent SNPs using PLINK1.9 (criteria linkage disequilibrium $r^{2}$=0.1 within windows of 10 Mbp). We calculated PGS for seven *P*-value thresholds (*P*<1.0E-05, *P*<0.001, *P*<0.01, *P*<0.05, *P*<0.1, *P*<0.5, and *P*<1.0) of the GWAS summary statistics. PGS were calculated from the imputed genotype dosages to the 1000 Genomes (Phase 3 Release 5) reference panel. We fitted linear mixed models, which controlled for relatedness using a genetic relatedness matrix (GRM) and covariates sex, age, two dummy variables for the GWAS array used, and the first five genetic PCs. The parameters of the model were estimated using GCTA 1.9 [68]. The linear model was as follows:

$$CD symptom score=intercept+Covariates*b+c*PGC+G$$

where $b$ and $c$ represent the vectors of fixed effects; and $G\sim N\left( 0,GRM*\sigma2G \right)$ represents the random effect that models the sample relatedness, with $GRM$ being the $N$ by $N$ matrix of relatedness estimated from SNPs, and $N$=10,706 is the number of individuals.

**Differential genetic correlation across rater-specific assessment of AGG and external outcomes**

We separately computed genetic correlations between rater-specific assessment of AGG and a list of external phenotypes (*N*=46). Note, since the GWAMA on paternal assessment of AGG returned a non-significant SNP-heritability ($h_{SNP}^{2}$=0.0412; SE=0.0261), rater-specific genetic correlations with external outcomes were only computed for maternal-, self-, and teacher-reported AGG. We applied Genomic Structural Equation Modelling (Genomic SEM; Grotzinger et al. 2019) to test whether genetic correlations between AGG and external phenotypes are significantly different across rater. Specifically, we considered the genetic correlations between the outcome and tree rater-specific assessments of AGG and fitted three models. In the first model, the genetic correlations between the outcome and all three rater-specific assessments of AGG were constrained at zero (null model). In the second model, the genetic correlations were allowed to differ from zero but constrained to be equal across raters (equal model). In the third model, the genetic correlations between rater-specific assessment of AGG and the outcome were freely estimated (free model). To facilitate model convergence, path loadings from the latent AGG variables on their respective observed variables, and genetic correlations between rater-specific assessment of AGG were constrained to be positive in all three models. Since the null model and equal model were nested inside the free model, model selection was performed based on a log-likelihood ratio test. To correct for multiple testing, we applied an FDR correction for 2 x 46 tests.

**References**

1.	Fraser A, Macdonald-Wallis C, Tilling K, Boyd A, Golding J, Davey Smith G, et al. Cohort Profile: The Avon Longitudinal Study of Parents and Children: ALSPAC mothers cohort. Int J Epidemiol. 2013;42:97–110.

2.	Boyd A, Golding J, Macleod J, Lawlor DA, Fraser A, Henderson J, et al. Cohort Profile: The ‘Children of the 90s’—the index offspring of the Avon Longitudinal Study of Parents and Children. Int J Epidemiol. 2013;42:111–127.

3.	Sunyer J, Esnaola M, Alvarez-Pedrerol M, Forns J, Rivas I, López-Vicente M, et al. Association between Traffic-Related Air Pollution in Schools and Cognitive Development in Primary School Children: A Prospective Cohort Study. PLOS Med. 2015;12:e1001792.

4.	Anckarsäter H, Lundström S, Kollberg L, Kerekes N, Palm C, Carlström E, et al. The Child and Adolescent Twin Study in Sweden (CATSS). Twin Res Hum Genet. 2011;14:495–508.

5.	Lichtenstein P, Tuvblad C, Larsson H, Carlström E. The Swedish Twin study of CHild and Adolescent Development: The TCHAD-Study. Twin Res Hum Genet. 2007;10:67–73.

6.	Brikell I, Larsson H, Lu Y, Pettersson E, Chen Q, Kuja-Halkola R, et al. The contribution of common genetic risk variants for ADHD to a general factor of childhood psychopathology. Mol Psychiatry. 2018:1–13.

7.	Fergusson DM, Horwood JL. The Christchurch Health and Development Study: Review of Findings on Child and Adolescent Mental Health. Aust New Zeal J Psychiatry. 2001;35:287–296.

8.	Fergusson DM, Horwood LJ. The Christchurch Health and Development Study. Christchurch Exp. 40 Years Res. Teach., Christchurch: University of Otago; 2013. p. 79–87.

9.	Adkins DE, Clark SL, Copeland WE, Kennedy M, Conway K, Angold A, et al. Genome-Wide Meta-Analysis of Longitudinal Alcohol Consumption Across Youth and Early Adulthood. Twin Res Hum Genet. 2015;18:335–347.

10.	Costello EJ, Eaves L, Sullivan P, Kennedy M, Conway K, Adkins DE, et al. Genes, Environments, and Developmental Research: Methods for a Multi-Site Study of Early Substance Abuse. Twin Res Hum Genet. 2013;16:505–515.

11.	Bisgaard H, Vissing NH, Carson CG, Bischoff AL, Følsgaard N V., Kreiner-Møller E, et al. Deep phenotyping of the unselected COPSAC _2010_ birth cohort study. Clin Exp Allergy. 2013;43:1384–1394.

12.	Poulton R, Moffitt TE, Silva PA. The Dunedin Multidisciplinary Health and Development Study: overview of the first 40 years, with an eye to the future. Soc Psychiatry Psychiatr Epidemiol. 2015;50:679–693.

13.	Moffitt TE. Teen-aged mothers in contemporary Britain. J Child Psychol Psychiatry. 2002;43:727–742.

14.	Kaprio J, Pulkkinen L, Rose RJ. Genetic and Environmental Factors in Health-related Behaviors: Studies on Finnish Twins and Twin Families. Twin Res. 2002;5:366–371.

15.	Kaprio J. Twin Studies in Finland 2006. Twin Res Hum Genet. 2006;9:772–777.

16.	Kaprio J. The Finnish Twin Cohort Study: An Update. Twin Res Hum Genet. 2013;16:157–162.

17.	Kooijman MN, Kruithof CJ, van Duijn CM, Duijts L, Franco OH, van IJzendoorn MH, et al. The Generation R Study: design and cohort update 2017. Eur J Epidemiol. 2016;31:1243–1264.

18.	Medina-Gomez C, Felix JF, Estrada K, Peters MJ, Herrera L, Kruithof CJ, et al. Challenges in conducting genome-wide association studies in highly admixed multi-ethnic populations: the Generation R Study. Eur J Epidemiol. 2015;30:317–330.

19.	Heinrich J, Bolte G, Hölscher B, Douwes J, Lehmann I, Fahlbusch B, et al. Allergens and endotoxin on mothers’ mattresses and total immunoglobulin E in cord blood of neonates. Eur Respir J. 2002;20:617–623.

20.	Berg A v., Krämer U, Link E, Bollrath C, Heinrich J, Brockow I, et al. Impact of early feeding on childhood eczema: Development after nutritional intervention compared with the natural course - The GINIplus study up to the age of 6 years. Clin Exp Allergy. 2010;40:627–636.

21.	Copeland WE, Angold A, Shanahan L, Costello EJ. Longitudinal Patterns of Anxiety From Childhood to Adulthood: The Great Smoky Mountains Study. J Am Acad Child Adolesc Psychiatry. 2014;53:21–33.

22.	Costello EJ, Angold A, Burns BJ, Stangl DK, Tweed DL, Erkanli A, et al. The Great Smoky Mountains Study of Youth. Arch Gen Psychiatry. 1996;53:1129.

23.	Rhea S-A, Bricker JB, Wadsworth SJ, Corley RP. The Colorado Adoption Project. Twin Res Hum Genet. 2013;16:358–365.

24.	Rhea S-A, Gross AA, Haberstick BC, Corley RP. Colorado Twin Registry: An Update. Twin Res Hum Genet. 2013;16:351–357.

25.	Derringer J, Corley RP, Haberstick BC, Young SE, Demmitt BA, Howrigan DP, et al. Genome-Wide Association Study of Behavioral Disinhibition in a Selected Adolescent Sample. Behav Genet. 2015;45:375–381.

26.	Guxens M, Ballester F, Espada M, Fernández MF, Grimalt JO, Ibarluzea J, et al. Cohort Profile: The INMA—INfancia y Medio Ambiente—(Environment and Childhood) Project. Int J Epidemiol. 2012;41:930–940.

27.	Iacono WG, Carlson SR, Taylor J, Elkins IJ, McGue M. Behavioral disinhibition and the development of substance-use disorders: Findings from the Minnesota Twin Family Study. Dev Psychopathol. 1999;11:869–900.

28.	Keyes MA, Malone SM, Elkins IJ, Legrand LN, McGue M, Iacono WG. The Enrichment Study of the Minnesota Twin Family Study: Increasing the Yield of Twin Families at High Risk for Externalizing Psychopathology. Twin Res Hum Genet. 2009;12:489–501.

29.	Magnus P, Birke C, Vejrup K, Haugan A, Alsaker E, Daltveit AK, et al. Cohort Profile Update: The Norwegian Mother and Child Cohort Study (MoBa). Int J Epidemiol. 2016;45:382–388.

30.	Burt SA, Klump KL. The Michigan State University Twin Registry (MSUTR): An Update. Twin Res Hum Genet. 2013;16:344–350.

31.	Keeping JD, Najman JM, Morrison J, Western JS, Andersen MJ, Williams GM. A prospective longitudinal study of social, psychological and obstetric factors in pregnancy: response rates and demographic characteristics of the 8556 respondents. BJOG An Int J Obstet Gynaecol. 1989;96:289–297.

32.	Najman JM, Alati R, Bor W, Clavarino A, Mamun A, McGrath JJ, et al. Cohort Profile Update: The Mater-University of Queensland Study of Pregnancy (MUSP). Int J Epidemiol. 2015;44:78-78f.

33.	Najman J, Bor W, O’Callaghan M, Williams G, Aird R, Shuttlewood G. Cohort Profile: The Mater-University of Queensland Study of Pregnancy (MUSP). Int J Epidemiol. 2005;34:992–997.

34.	Järvelin MR, Hartikainen-Sorri A-L, Rantakallio P. Labour induction policy in hospitals of different levels of specialisation. BJOG An Int J Obstet Gynaecol. 1993;100:310–315.

35.	van Beijsterveldt CEM, Groen-Blokhuis M, Hottenga JJ, Franić S, Hudziak JJ, Lamb D, et al. The Young Netherlands Twin Register (YNTR): Longitudinal Twin and Family Studies in Over 70,000 Children. Twin Res Hum Genet. 2013;16:252–267.

36.	Bartels M, Van Beijsterveldt CEM, Derks EM, Stroet TM, Polderman TJC, Hudziak JJ, et al. Young Netherlands Twin Register (Y-NTR): A longitudinal multiple informant study of problem behavior. Twin Res Hum Genet. 2007;10:3–11.

37.	Hudziak JJ, van Beijsterveldt CEM, Bartels M, Rietveld MJH, Rettew DC, Derks EM, et al. Individual Differences in Aggression: Genetic Analyses by Age, Gender, and Informant in 3-, 7-, and 10-Year-Old Dutch Twins. Behav Genet. 2003;33:575–589.

38.	Wesseldijk LW, Bartels M, Vink JM, van Beijsterveldt CEM, Ligthart L, Boomsma DI, et al. Genetic and environmental influences on conduct and antisocial personality problems in childhood, adolescence, and adulthood. Eur Child Adolesc Psychiatry. 2018;27:1123–1132.

39.	Scheet P, Ehli EA, Xiao X, van Beijsterveldt CEM, Abdellaoui A, Althoff RR, et al. Twins, Tissue, and Time: An Assessment of SNPs and CNVs. Twin Res Hum Genet. 2012;15:737–745.

40.	Heath AC, Whitfield JB, Martin NG, Pergadia ML, Goate AM, Lind PA, et al. A Quantitative-Trait Genome-Wide Association Study of Alcoholism Risk in the Community: Findings and Implications. Biol Psychiatry. 2011;70:513–518.

41.	KNOPIK VS, HEATH AC, MADDEN PAF, BUCHOLZ KK, SLUTSKE WS, NELSON EC, et al. Genetic effects on alcohol dependence risk: re-evaluating the importance of psychiatric and other heritable risk factors. Psychol Med. 2004;34:1519–1530.

42.	Medland SE, Nyholt DR, Painter JN, McEvoy BP, McRae AF, Zhu G, et al. Common Variants in the Trichohyalin Gene Are Associated with Straight Hair in Europeans. Am J Hum Genet. 2009;85:750–755.

43.	Newnham JP, Evans SF, Michael CA, Stanley FJ, Landau LI. Effects of frequent ultrasound during pregnancy: a randomised controlled trial. Lancet. 1993;342:887–891.

44.	Straker L, Mountain J, Jacques A, White S, Smith A, Landau L, et al. Cohort Profile: The Western Australian Pregnancy Cohort (Raine) Study–Generation 2. Int J Epidemiol. 2017;46:dyw308.

45.	Haworth CMA, Davis OSP, Plomin R. Twins Early Development Study (TEDS): A Genetically Sensitive Investigation of Cognitive and Behavioral Development From Childhood to Young Adulthood. Twin Res Hum Genet. 2013;16:117–125.

46.	de Winter AF, Oldehinkel AJ, Veenstra R, Brunnekreef JA, Verhulst FC, Ormel J. Evaluation of non-response bias in mental health determinants and outcomes in a large sample of pre-adolescents. Eur J Epidemiol. 2005;20:173–181.

47.	Oldehinkel AJ, Rosmalen JG, Buitelaar JK, Hoek HW, Ormel J, Raven D, et al. Cohort Profile Update: The TRacking Adolescents’ Individual Lives Survey (TRAILS). Int J Epidemiol. 2015;44:76-76n.

48.	Simonoff E, Pickles A, Meyer JM, Silberg JL, Maes HH, Loeber R, et al. The Virginia Twin Study of Adolescent Behavioral Development. Arch Gen Psychiatry. 1997;54:801.

49.	Hewitt JK, Rutter M, Simonoff E, Pickles A, Loeber R, Heath AC, et al. Genetics and Developmental Psychopathology: 1. Phenotypic Assessment in the Virginia Twin Study of Adolescent Behavioral Development. J Child Psychol Psychiatry. 1997;38:943–963.

50.	Eaves LJ, Silberg JL, Meyer JM, Maes HH, Simonoff E, Pickles A, et al. Genetics and Developmental Psychopathology: 2. The Main Effects of Genes and Environment on Behavioral Problems in the Virginia Twin Study of Adolescent Behavioral Development. J Child Psychol Psychiatry. 1997;38:965–980.

51.	Maes HH, Silberg JL, Neale MC, Eaves LJ. Genetic and Cultural Transmission of Antisocial Behavior: An Extended Twin Parent Model. Twin Res Hum Genet. 2007;10:136–150.

52.	Raitakari OT, Juonala M, Ronnemaa T, Keltikangas-Jarvinen L, Rasanen L, Pietikainen M, et al. Cohort Profile: The Cardiovascular Risk in Young Finns Study. Int J Epidemiol. 2008;37:1220–1226.

53.	Juonala M, Viikari JSA, Raitakari OT. Main findings from the prospective Cardiovascular Risk in Young Finns Study. Curr Opin Lipidol. 2013;24:57–64.

54.	Boomsma DI. Aggression in children: unravelling the interplay of genes and environment through (epi)genetics and metabolomics. J Pediatr Neonatal Individ Med. 2015;4:e040251.

55.	Bartels M, Hendriks A, Mauri M, Krapohl E, Whipp A, Bolhuis K, et al. Childhood aggression and the co-occurrence of behavioural and emotional problems: results across ages 3–16 years from multiple raters in six cohorts in the EU-ACTION project. Eur Child Adolesc Psychiatry. 2018;27:1105–1121.

56.	Middeldorp CM, Felix JF, Mahajan A, McCarthy MI, consortium EGG (EGG), McCarthy MI. The Early Growth Genetics (EGG) and EArly Genetics and Lifecourse Epidemiology (EAGLE) consortia: design, results and future prospects. Eur J Epidemiol. 2019:1–22.

57.	Achenbach TM, Ivanova MY, Rescorla LA. Empirically based assessment and taxonomy of psychopathology for ages 1½–90+ years: Developmental, multi-informant, and multicultural findings. Compr Psychiatry. 2017;79:4–18.

58.	Goodman R. Psychometric Properties of the Strengths and Difficulties Questionnaire. J Am Acad Child Adolesc Psychiatry. 2001;40:1337–1345.

59.	Minică CC, Dolan C V, Kampert MMD, Boomsma DI, Vink JM. Sandwich corrected standard errors in family-based genome-wide association studies. Eur J Hum Genet. 2015;23:388–394.

60.	Winkler TW, Day FR, Croteau-Chonka DC, Wood AR, Locke AE, Mägi R, et al. Quality control and conduct of genome-wide association meta-analyses. Nat Protoc. 2014;9:1192–1212.

61.	Liu Q, Cirulli ET, Han Y, Yao S, Liu S, Zhu Q. Systematic assessment of imputation performance using the 1000 Genomes reference panels. Brief Bioinform. 2015;16:549–562.

62.	Bulik-Sullivan B, Finucane HK, Anttila V, Gusev A, Day FR, Loh P-R, et al. An atlas of genetic correlations across human diseases and traits. Nat Genet. 2015;47:1236–1241.

63.	Baselmans BML, Jansen R, Ip HF, van Dongen J, Abdellaoui A, van de Weijer MP, et al. Multivariate genome-wide analyses of the well-being spectrum. Nat Genet. 2019:1.

64.	Bulik-Sullivan BK, Loh P-R, Finucane HK, Ripke S, Yang J, Schizophrenia Working Group of the Psychiatric Genomics Consortium, et al. LD Score regression distinguishes confounding from polygenicity in genome-wide association studies. Nat Genet. 2015;47:291–295.

65.	Chang CC, Chow CC, Tellier LC, Vattikuti S, Purcell SM, Lee JJ. Second-generation PLINK: rising to the challenge of larger and richer datasets. Gigascience. 2015;4:7.

66.	R Core Team. R: A language and environment for statistical computing. 2019.

67.	Rogers P, Stoner J. Modification of the Sandwich Estimator in Generalized Estimating Equations with Correlated Binary Outcomes in Rare Event and Small Sample Settings. Am J Appl Math Stat. 2015;3:243–251.

68.	Yang J, Lee SH, Goddard ME, Visscher PM. GCTA: a tool for genome-wide complex trait analysis. Am J Hum Genet. 2011;88:76–82.

69.	Grotzinger AD, Rhemtulla M, de Vlaming R, Ritchie SJ, Mallard TT, Hill WD, et al. Genomic structural equation modelling provides insights into the multivariate genetic architecture of complex traits. Nat Hum Behav. 2019:1.

**Supplementary Figures**

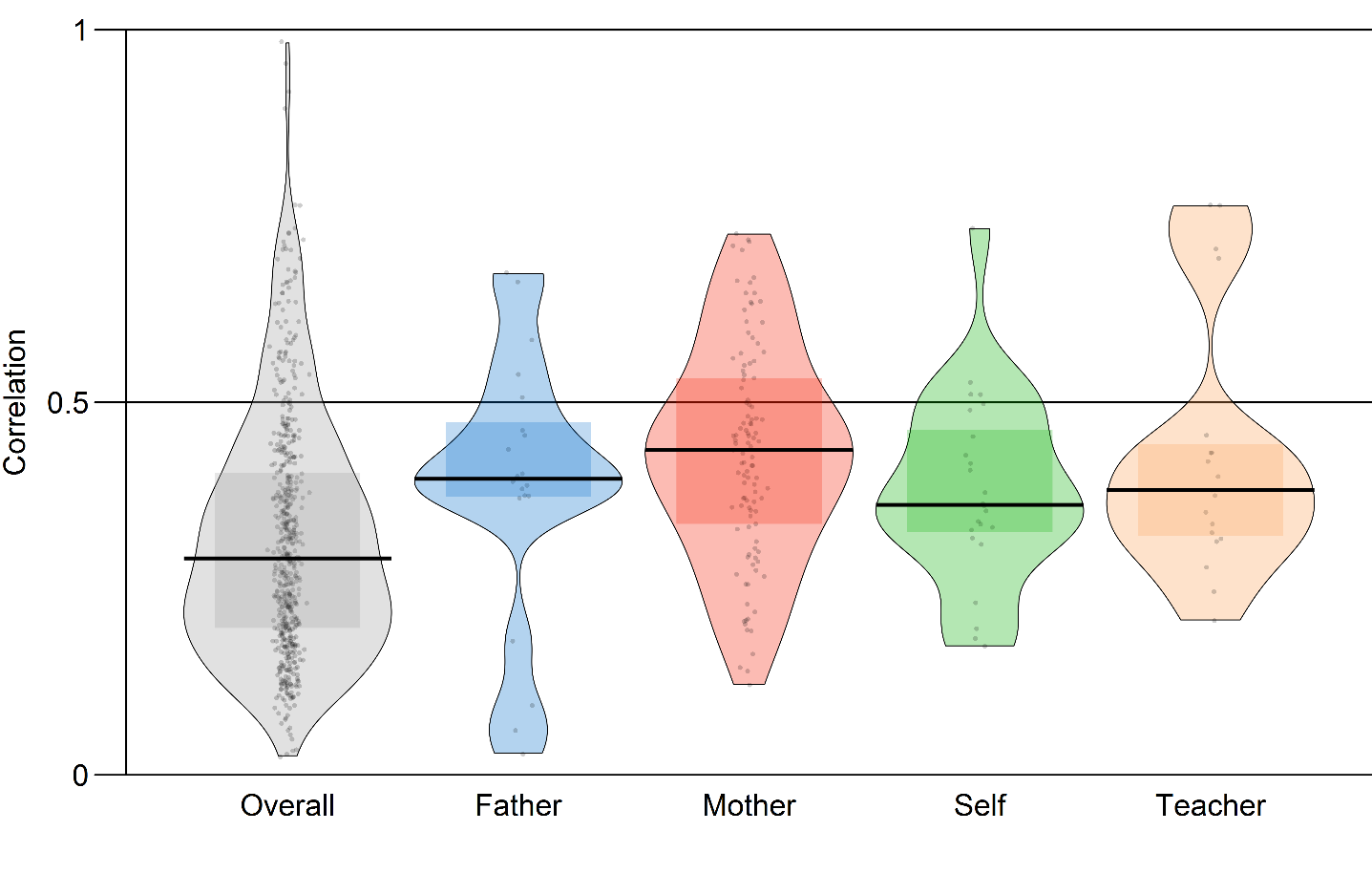

**Supplementary Figure 1.** Distribution of phenotypic correlations. From left to right, colors represent (1) correlations across all samples, (2) correlations within father-reported data, (3) mother-reported data, (4) self-reported data, and (5) teacher-reported data. Boxes represent the interquartile range.

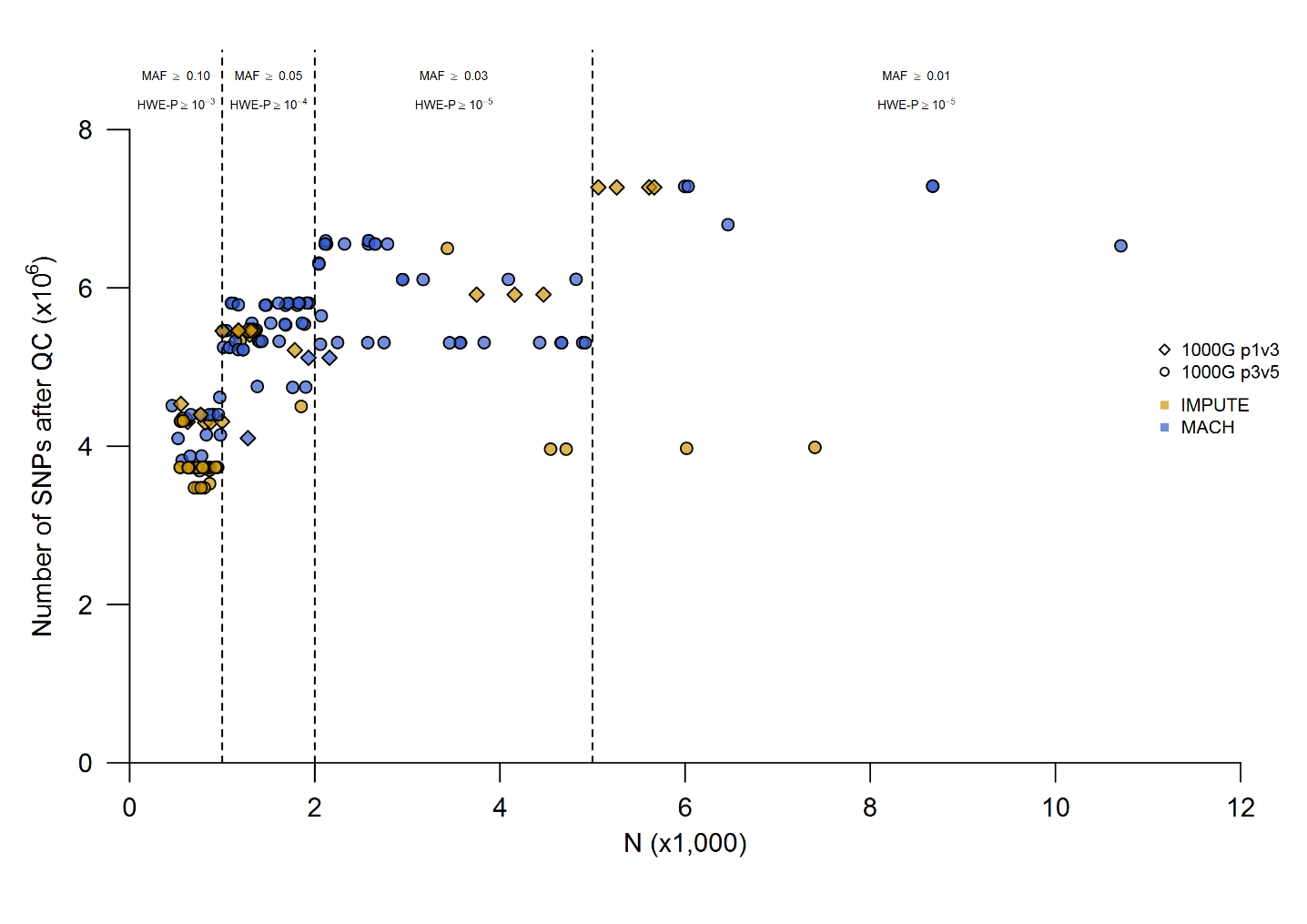

**Supplementary Figure 2.** Number of SNPs after QC per GWAS. Horizontal axis shows the (maximum) sample size. Vertical axis shows the number of SNPs after QC. Shape of the points correspond to imputation reference panel. Color of the points correspond to software used for imputation. Dotted vertical lines represent the upper limit of the sample-size-dependent QC filters. MAF=minor allele frequency; HWE-P=Hardy-Weinberg equilibrium test *P*-value.

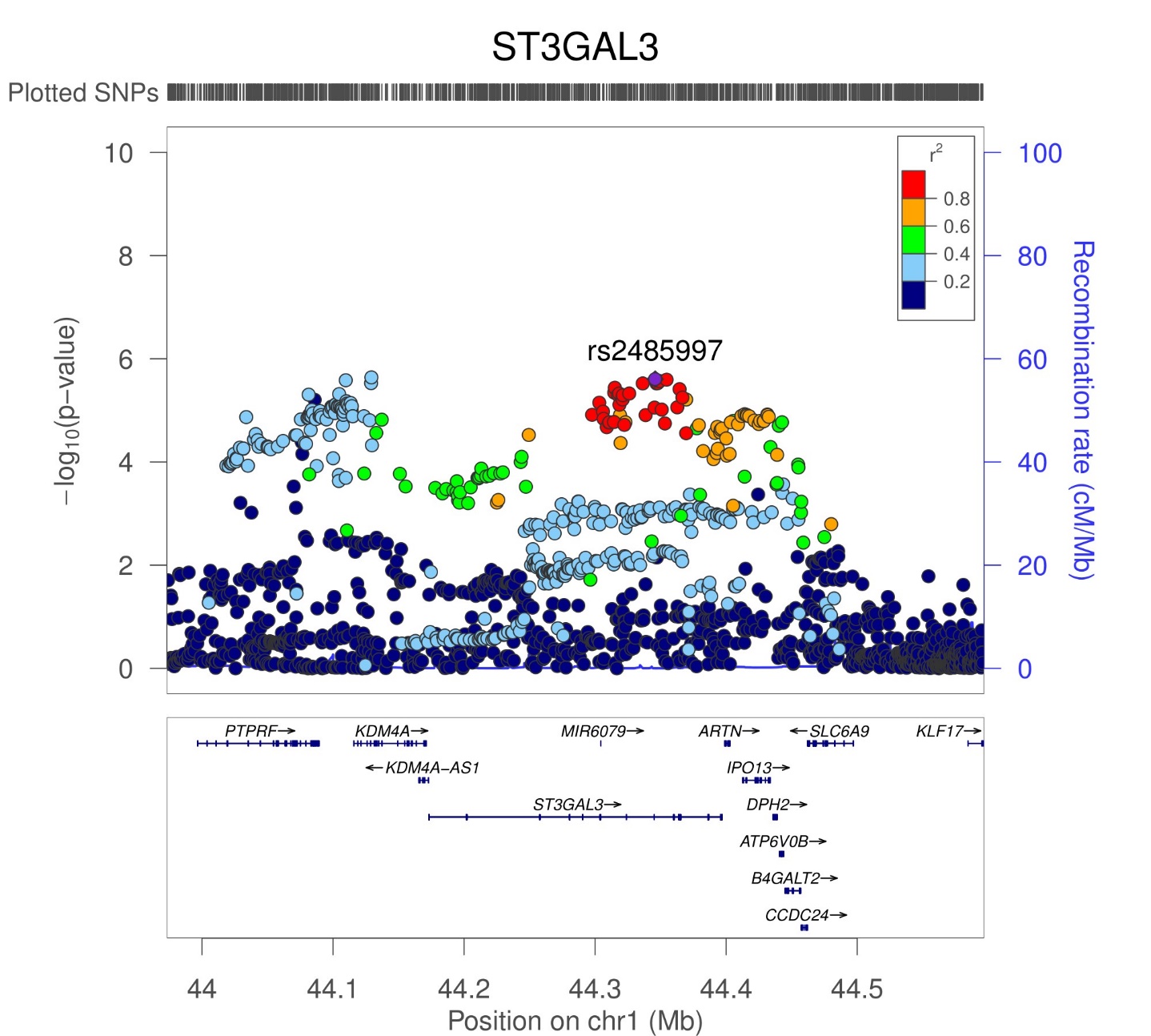

**Supplementary Figure 3.** Zoom plot of *ST3GAL3*. Plotting window was extended with 200Kbp on both sides of the gene. Purple diamond indicates the index variant inside the gene.

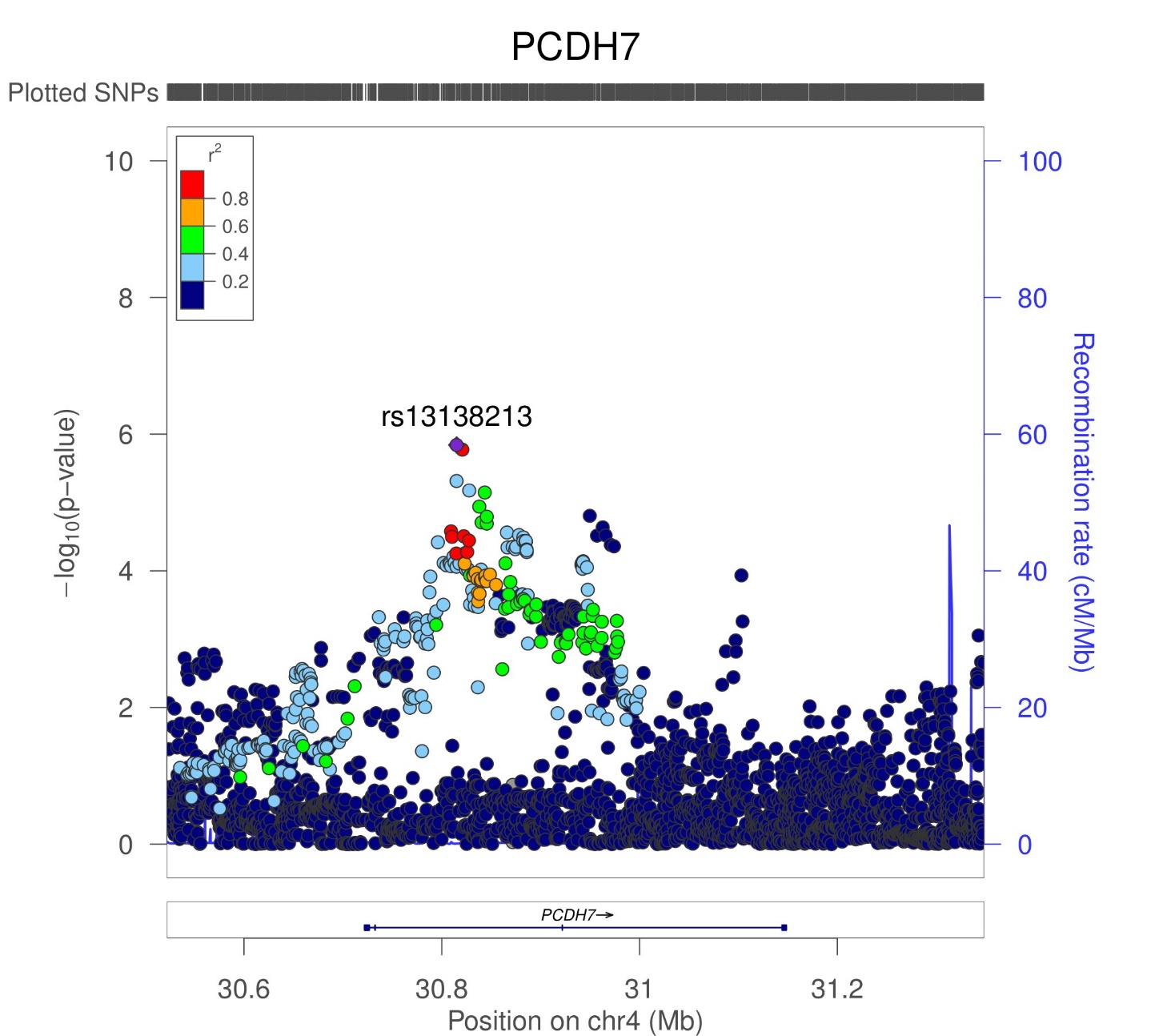

**Supplementary Figure 4.** Zoom plot of *PCDH7*. Plotting window was extended with 200Kbp on both sides of the gene. Purple diamond indicates the index variant inside the gene.

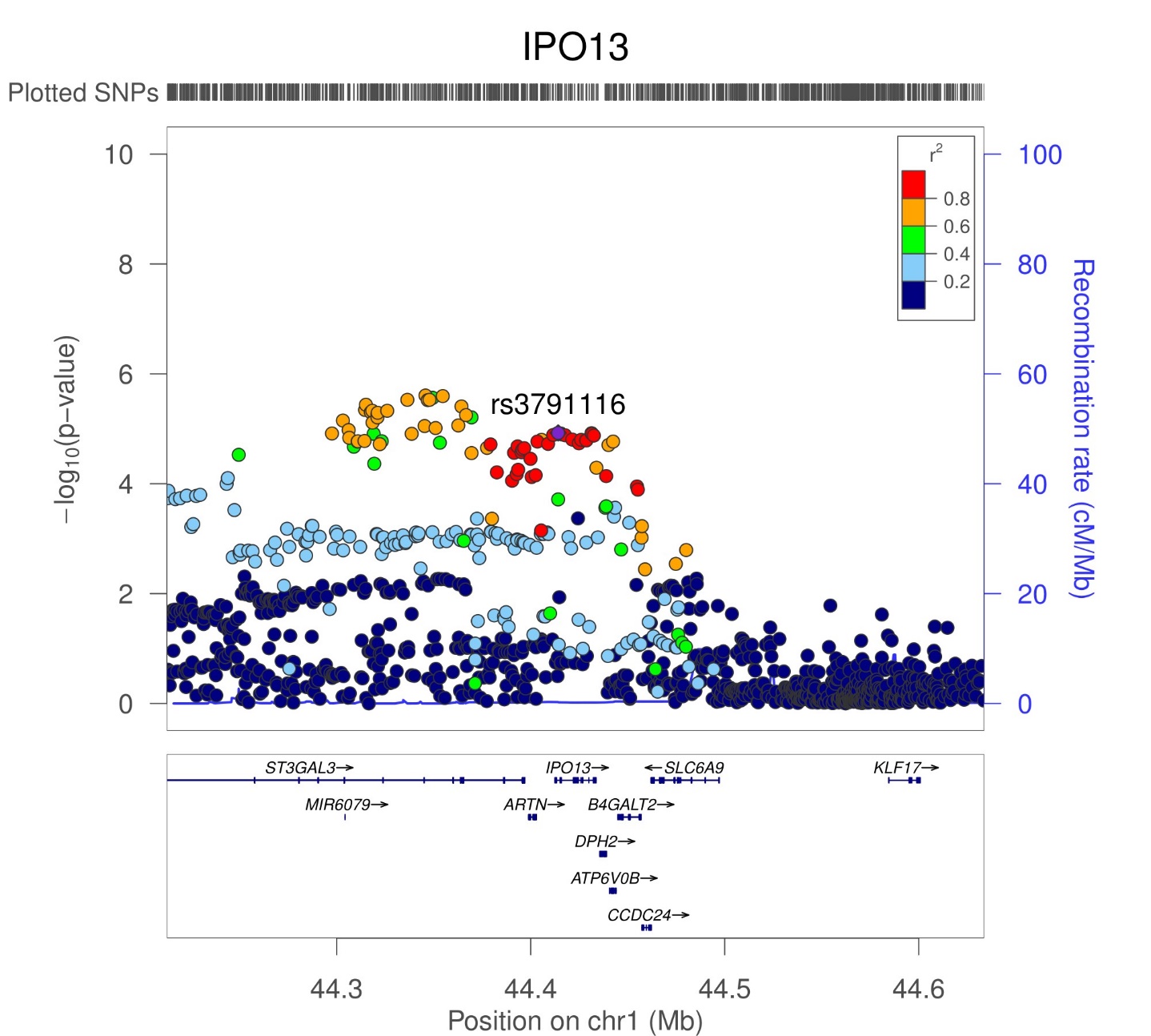

**Supplementary Figure 5.** Zoom plot of *IPO13*. Plotting window was extended with 200Kbp on both sides of the gene. Purple diamond indicates the index variant inside the gene.

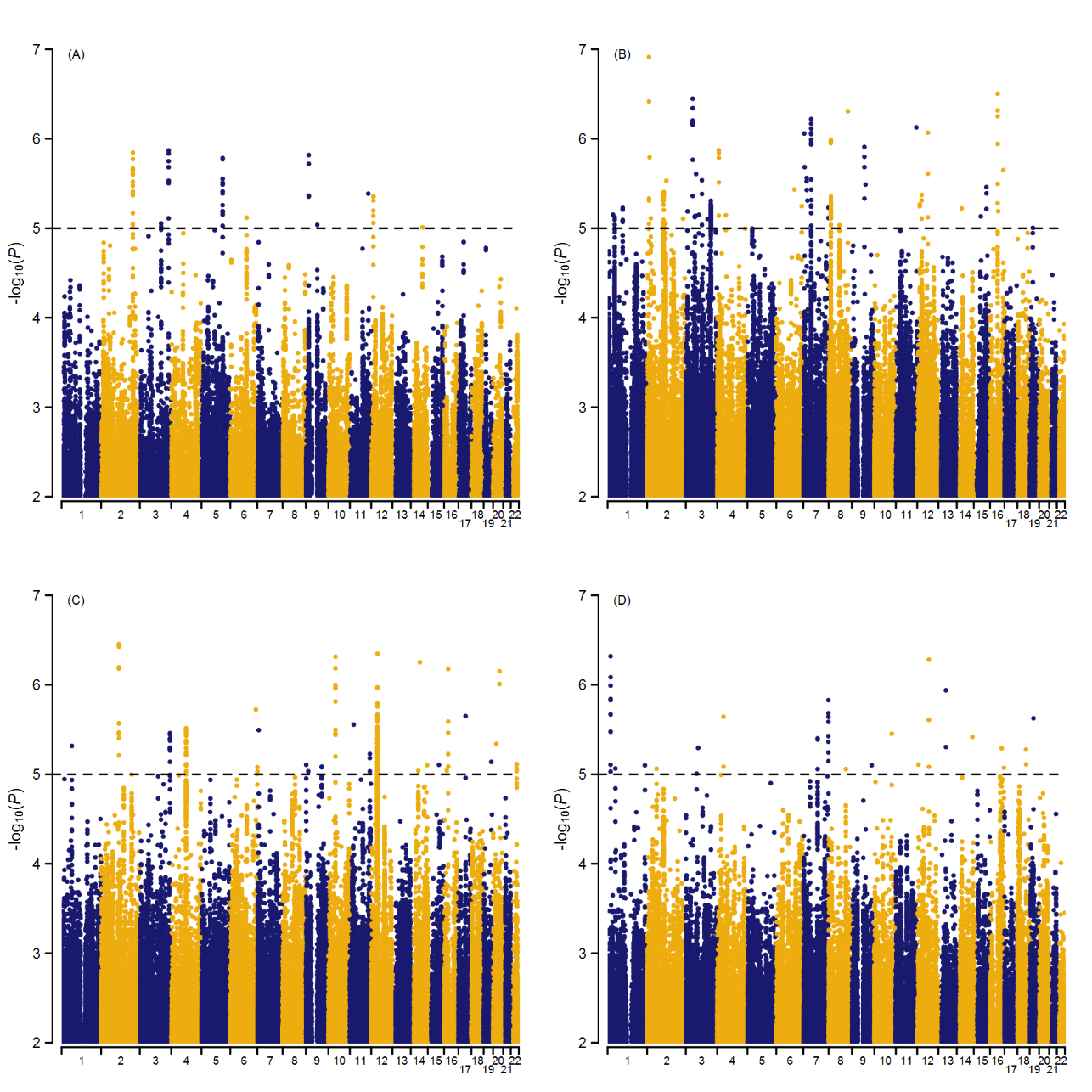

**Supplementary Figure 6.** Manhattan plot per rater-specific GWAMA. (A) father. (B) mother. (C) self. (D) teacher. Dotted horizontal line represents threshold for suggestive association (*P*=10^-5^).

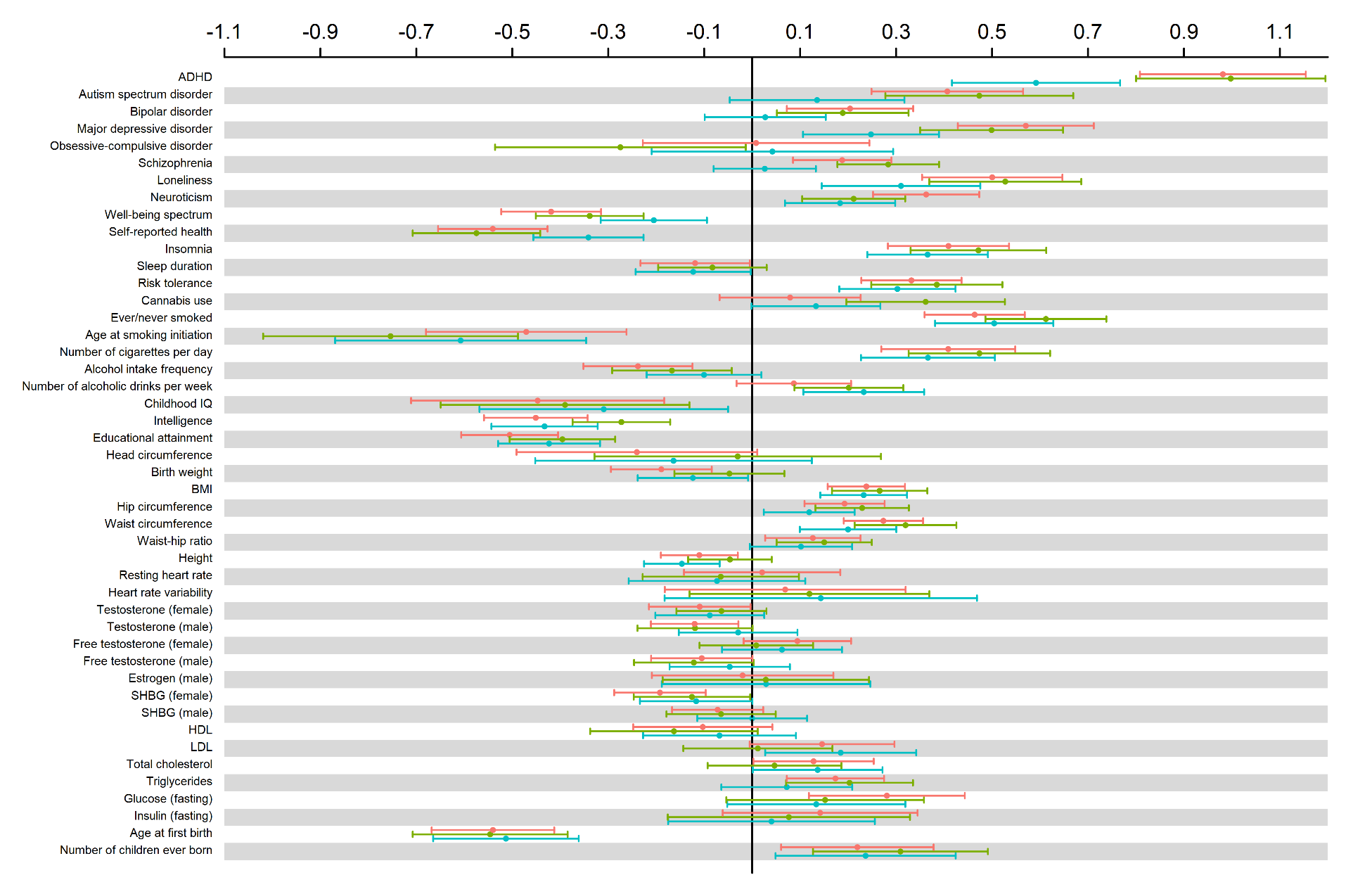

**Supplementary Figure 7**. Genetic correlations between rater-specific assessment of AGG and external phenotypes. Phenotypes are sorted on domain. Colors represent raters: red=mother, green=self, blue=teacher. Bars indicate 95% confidence intervals.
